# Excitable dynamics of Ras triggers self-organized PIP3 signaling for spontaneous cell migration

**DOI:** 10.1101/356105

**Authors:** Seiya Fukushima, Satomi Matsuoka, Masahiro Ueda

**Affiliations:** Department of Biological Science, Graduate School of Science, Osaka University, Toyonaka, Osaka 560-0043, Japan; RIKEN Center for Biosystems Dynamics Research (BDR), Suita, Osaka 565-0874, Japan; Graduate School of Frontier Biosciences, Osaka University, Suita, Osaka 565-0871, Japan

## Abstract

Spontaneous cell movement is underpinned by an asymmetric distribution of signaling molecules including small G proteins and phosphoinositides on the cell membrane. A fundamental question is the molecular mechanism for the spontaneous symmetry breaking. Here we report that GTP bound Ras (Ras-GTP) breaks the symmetry due to excitability even in the absence of extracellular spatial cues and cytoskeletal polarity as well as downstream signaling activities. A stochastic excitation of local and transient Ras activation induced PIP3 accumulation via direct interaction with PI3K, causing tightly coupled traveling waves propagating along the membrane. Comprehensive phase analysis of the waves of Ras-GTP and PIP3 metabolism-related molecules revealed the network structure of the excitable system including positive feedback regulation of Ras-GTP by PIP3. A mathematical model reconstituted a series of the observed symmetry breaking phenomena, illustrating an essential involvement of excitability in the cellular decision-making process.

**Author contributions:** S.F. conducted the experiments; all authors analyzed the data and wrote the manuscript.

**Abbreviations**
GAP: Guanosine Triphosphate hydrolase activating protein
GEF: Guanine nucleotide exchange factor
GFP: Green fluorescent protein
PHD: Pleckstrin homology domain
PI3K: Phosphoinositide 3-kinase
PIP2: Phosphatidylinositol (4,5)-bisphosphate
PIP3: Phosphatidylinositol (3,4,5)-trisphosphate
PKB&Akt: Protein kinase B
PLCδ1: Phospholipase C delta 1
PTEN: Phosphatase and tensin homolog
RBD: Ras binding domain
RFP: Red fluorescent protein

## Introduction

Symmetry breaking underlies essential cellular decision-making processes including polarization, migration and division. In eukaryotic amoeboid cells, intracellular asymmetric signals generate cell motility through cytoskeletal rearrangement [1,2]. Several studies have investigated the underpinning signaling mechanisms in various chemotactic cells and found common features in the network structures [3–7]. For example, multiple signaling molecules including phosphatidylinositol lipids, Ras GTPases and various kinases accumulate locally on the cell periphery [8,9]. These asymmetric signals occur even in the absence of functional actin cytoskeleton or external asymmetry in chemoattractant stimulations [10–13]. Therefore, the signaling network itself has the ability to exhibit symmetry breaking as internal spontaneous dynamics. However, it remains to be fully elucidated how the asymmetric signal emerges from the intracellular signaling network.

Recent evidence demonstrates that asymmetric signals can spontaneously arise from excitability of the signaling network [4,14,15]. Excitability in biology has been well documented experimentally and theoretically in the action potentials of neurons [16,17]. Similar findings have since been made in many other cellular phenomena such as cell differentiation [18,19], gene expression [20,21] and eukaryotic chemotaxis [3,22]. In general, an excitable system has a threshold for all-or-none excitation, ensuring that cells can respond only to a supra-threshold stimulus in an all-or-none manner. In addition, excitable systems can exhibit spontaneous excitation due to internal fluctuations or molecular noise without any external stimulus as a basis for the self-organization of oscillations or travelling waves [23]. These properties of excitable systems provide mechanisms for intracellular pattern formation, by which a signaling domain is generated locally on the cell membrane, leading to symmetry breaking.

Chemotactic signaling pathways in *Dictyostelium discoideum* exhibit the characteristics of excitable systems [5,14,15]. The phosphatidylinositol 3,4,5-trisphosphate (PIP3), TorC2, PLA2 and sGC pathways located downstream of chemoattractant receptors in parallel can each generate an intracellular cue for symmetry breaking in cell migration [10,11,24–27]. The chemoattractant gradient signals are mediated by G-protein-coupled receptors, heterotrimeric G proteins and Ras GTPases and bias the asymmetric signals along the gradient direction for chemotaxis [9]. In the PIP3 pathway, the PIP3-enriched domain acts as the asymmetric signal on the cell membrane at the front [4,28]. Evidence for excitability in the PIP3 pathway includes stimulation-induced all-or-none excitation, refractory behavior, spontaneous excitation and travelling wave generation [14,15,29,30]. Mammalian cells also exhibit the travelling wave-like localization patterns of PIP3 [31], implying an evolutionary conserved mechanism involving excitability as a common basis for the asymmetric signal generation in various chemotactic cells. However, which molecules in the signaling network work as key determinants for the emergence of excitable dynamics remains unclear.

Here, we performed quantitative live-cell imaging analysis to reveal the spatiotemporal relationship between several major signaling components including Ras-GTP and the PIP3 pathway in excitable dynamics. We found the excitability of Ras-GTP generates traveling waves (Ras waves) independently of downstream pathways to govern the self-organization of PIP3 traveling waves (PIP3 waves) via Ras-GTP-PI3K interactions. Based on the experimental results, we developed a reaction-diffusion model that reproduced the travelling waves, illustrating Ras is central to the emergence of excitable dynamics for asymmetric signal generation. Feedback regulation of the Ras excitability stabilized the asymmetric signal, suggesting the possible involvement of Ras-GTP in the signal integration for cell motility.

## Results

### Ras wave formation is independent of PIP3 and other downstream pathways

We performed live-cell imaging analysis of both Ras-GTP and PIP3 by using RBD_Raf1_-GFP (or RFP) and PHD_AKT/PKB_-GFP, two fluorescent reporters specific for Ras-GTP and PIP3, respectively [32]. Under confocal microscope observation, Ras-GTP and PIP3 exhibited travelling waves along the cell periphery in *Dictyostelium* cells (Fig 1A, S1 Movie), consistent with previous observations [14,30]. To avoid the effects of the actin cytoskeleton in the Ras-GTP and PIP3 dynamics, the cells were treated with an actin-polymerization inhibitor, latrunculin A. Following the method described previously [13], the cells were also treated with 4 mM caffeine to observe travelling waves along the membrane. A kymograph showing the intensities of both probes along the membrane indicated clearly co-localizing Ras and PIP3 waves in the background of wild type (WT) cells (Fig 1B).

**Figure 1.**
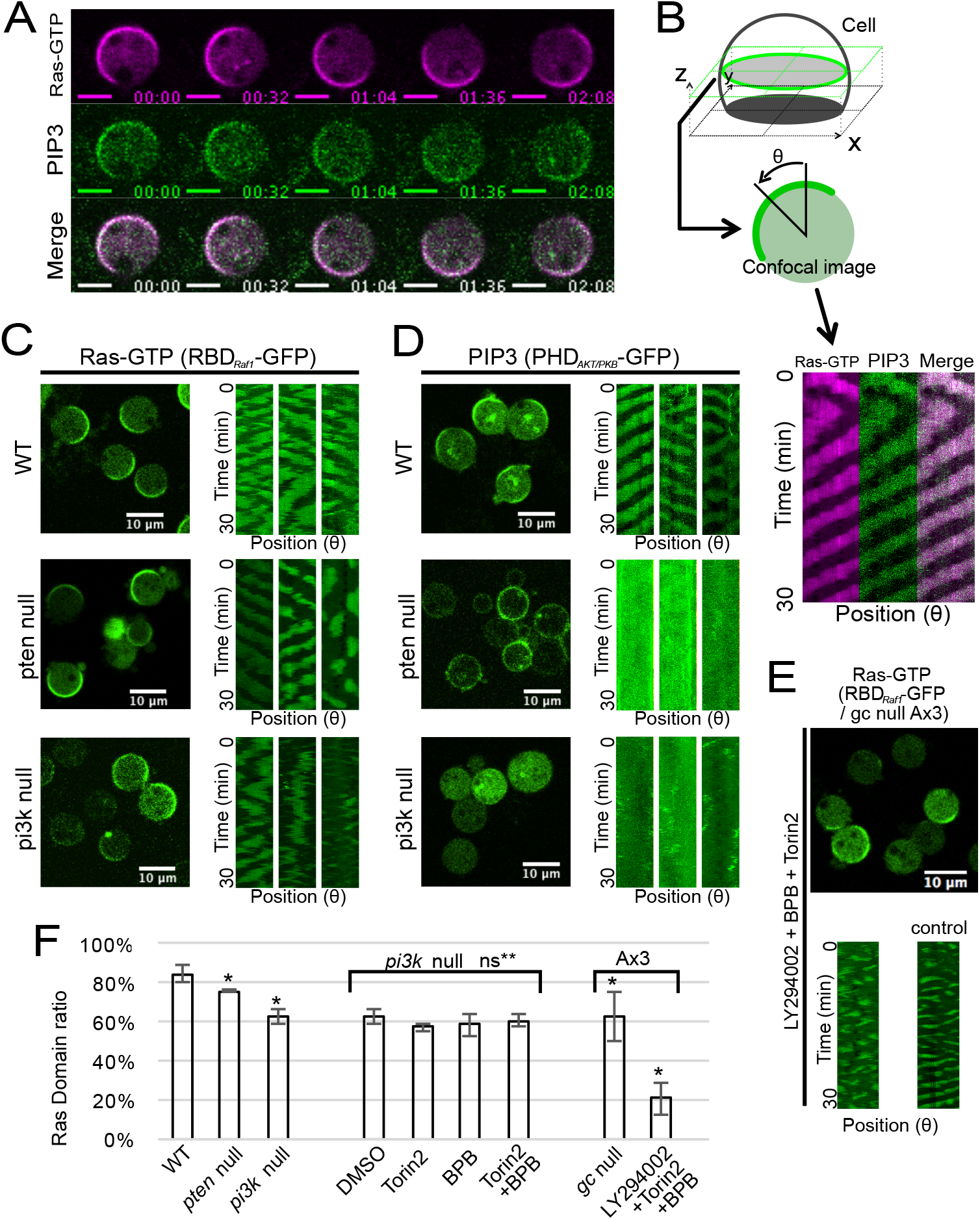
Ras waves in the absence of active downstream parallel pathways. (A) Simultaneous time-lapse imaging of Ras-GTP and PIP3 waves in WT cells expressing RBD Raf1-RFP and PHD_AKT/PKB_-GFP taken by confocal microscopy. Scale bars, 5 *μ*m. Time format is “mm:ss”. (B) Kymograph analysis. (C), (D) Confocal images and kymographs of Ras or PIP3 waves in WT, *pten* null and *pi3k1-5* null cells. (E) Confocal image and kymographs of *gc* null cells (Ax3 strain) treated with a combination of 100 *μ*M LY294002, 10 *μ*M Torin2 and 2 *μ*M BPB. **F,** Ratio of the number of cells showing Ras-GTP enriched domains. Data are mean ± s.d. of three independent experiments, and more than 200 cells were counted in each experiment. (* P < 0.01 Welch’s t-test against WT; ns** P > 0.01 Welch’s t-test against *pi3k1-5* null).

To see whether the traveling wave generation of Ras-GTP requires the PIP3 wave or not, we observed both probes in *pi3k1-5* null and *pten* null strains, because PI3K and PTEN are essential for the production and degradation of PIP3, respectively [27,33,34]. Ras-GTP exhibited wave propagation even without PI3K or PTEN, whereas PIP3 did not (Fig 1C and 1D). The efficiencies of Ras wave generation in WT, *pi3k1-5* and *pten* null cells were 84% (n=1541), 63% (n=1487) and 76% (n=1982), respectively (Fig 1F), indicating PIP3 production and degradation are not necessary for Ras wave generation. We noticed that Ras-GTP waves exhibited zigzag or disconnected patterns with prolonged oscillatory periods in both mutants compared with WT (S1C Fig), suggesting a partial contribution of PIP3 production and degradation to the maintenance of Ras waves.

We further examined the possible involvement of other parallel chemotactic signaling pathways in the Ras wave generation. When we applied Torin2 and BPB, which are inhibitors for the TorC2 and PLA2 pathways, respectively, to *pi3k1-5* null cells, no obvious changes in Ras waves were observed (Fig 1F, S1B Fig). Even in the cells with all major four pathways (the PIP3, TorC2, PLA2, and sGC pathways) inhibited, Ras waves were still observed but with some defects (Fig 1E and 1F, S1A Fig)[24,35]. Thus, none of the four major pathways that mediate chemotactic signals downstream of Ras-GTP were necessary for Ras wave formation. That is, the spatiotemporal dynamics of Ras GTPase is excitable without the activities of the downstream pathways.

### Ras waves trigger PIP3 waves via Ras-GTP-PI3K interaction

We examined temporal relationships among Ras-GTP, PI3K and PIP3 during the formation of self-organized patterns by using TIRFM (total internal reflection fluorescence microscopy). TIRFM can illuminate only the membrane, has low background noise and provides larger data sets with higher temporal resolution than confocal microscopy. Simultaneous observations of RBD_Raf1_-RFP and PHD_AKT/PKB_-GFP revealed that both probes exhibited closely coupled wave propagation on the membrane (Fig 2A, S2 Movie). We obtained time trajectories of the fluorescence intensities in a ROI and found that the PIP3 wave propagation was slightly delayed compared with the Ras wave propagation (Fig 2B and 2C), which was obvious in the average dynamics obtained from 90 individual trajectories from 15 cells (Fig 2D). We calculated the peak time of the crosscorrelation function in each cell and obtained the distribution (Fig 2G, S2C Fig), which showed the lag time of PIP3 against Ras-GTP was about 0.9 ± 0.6 s on average.

**Figure 2.**
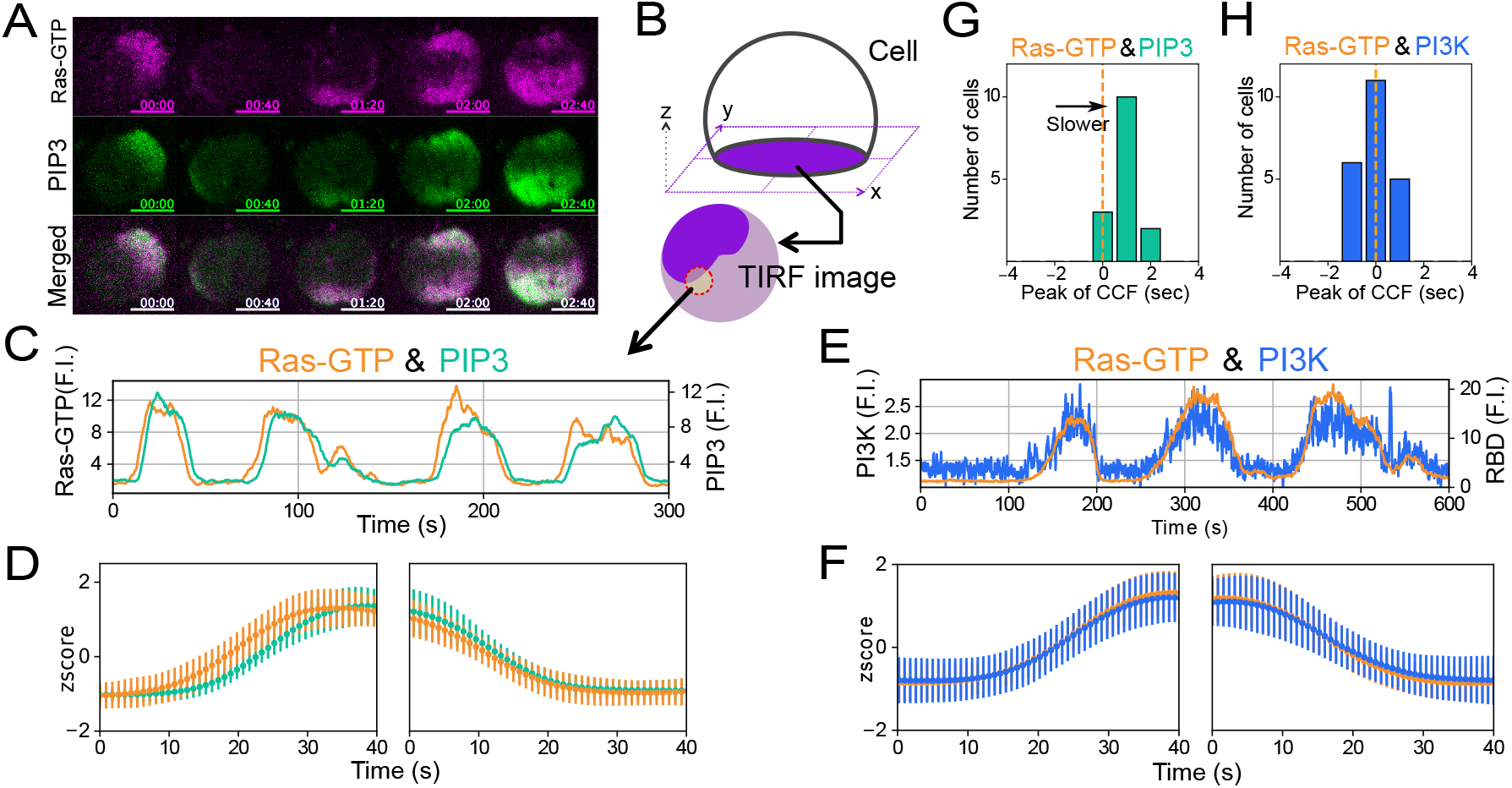
Synchronized propagation of Ras/PIP3/PI3K waves. (A) Simultaneous time-lapse imaging of Ras-GTP and PIP3 taken by TIRFM. Scale bars, 5 *μ*m. Time format is “mm:ss”. (B) Time trajectory analysis. (C), (E) Typical examples of time trajectories of Ras-GTP (orange), PIP3 (green) and PI3K (blue). Fluorescence intensity was normalized to the minimum in each trajectory. (D), (F) Average dynamics of the increasing (left) and decreasing phase (right). Orange, green and blue lines indicate Ras-GTP, PIP3 and PI3K, respectively. Data are the mean ± s.d. from more than 20 cells. (G), (H) Distribution of peak times of the cross-correlation functions. Dotted lines indicate time zero. The average peak value of Ras-GTP against PIP3 is 0.9 ± 0.6 s, against PI3K is 0.0 ± 0.7 s.

We next analyzed the temporal relationship between Ras-GTP and PI3K in the wave propagation. We focused on PI3K2, which makes a significant contribution to the catalytic activity for PIP3 production and cell migration, as a major effector of activated RasG among the six PI3K in *Dictyostelium discoideum* [11,36]. PI3K2 was labeled with tetramethylrhodamine (TMR) through HaloTag (PI3K2-Halo-TMR). Using TIRFM, we successfully visualized the wave dynamics of PI3K2-Halo-TMR on the membrane (S2A Fig). The oscillatory dynamics of PI3K2 on the cell membrane coincided tightly with that of Ras-GTP (Fig 2E and 2F, S2B Fig, S3 Movie). The peak time of the cross-correlation function was 0.0 ± 0.7 s on average (Fig 2H, S2D Fig), indicating no delay between the Ras-GTP and PI3K2 waves. When PI3K2-Halo-TMR and PHD_AKT/PKB_-GFP were observed simultaneously (S3 Fig, S4 Movie), the lag time of PIP3 against PI3K2 was about 2.3 ± 1.1 s on average, confirming the PIP3 waves follow the Ras/PI3K2 waves.

To confirm that the membrane localization of PI3K2 is regulated by interaction with Ras-GTP, we used a PI3K2 mutant, PI3K2^K857, 858E^, that is defective in Ras binding [11]. PI3K2^K857, 858E^ was distributed uniformly and slightly on the membrane with no wave generation during the Ras wave propagation in WT cells (Fig 3A and 3B, S5 Movie), indicating that the PI3K2 wave depends on the interaction with Ras-GTP. Furthermore, we confirmed that Ras-GTP/PI3K2 interaction is required for PIP3 wave generation by the rescue experiment. The *pi3k1-5* null cells exhibited PIP3 localization when transformed with PI3K2-Halo but not with PI3K2^K857, 858E^-Halo (Fig 3C, S2E Fig). These results together demonstrate that Ras-GTP excitation triggers PIP3 waves through Ras-GTP interaction with PI3K, and that PIP3 subordinates with Ras-GTP/PI3K.

**Figure 3.**
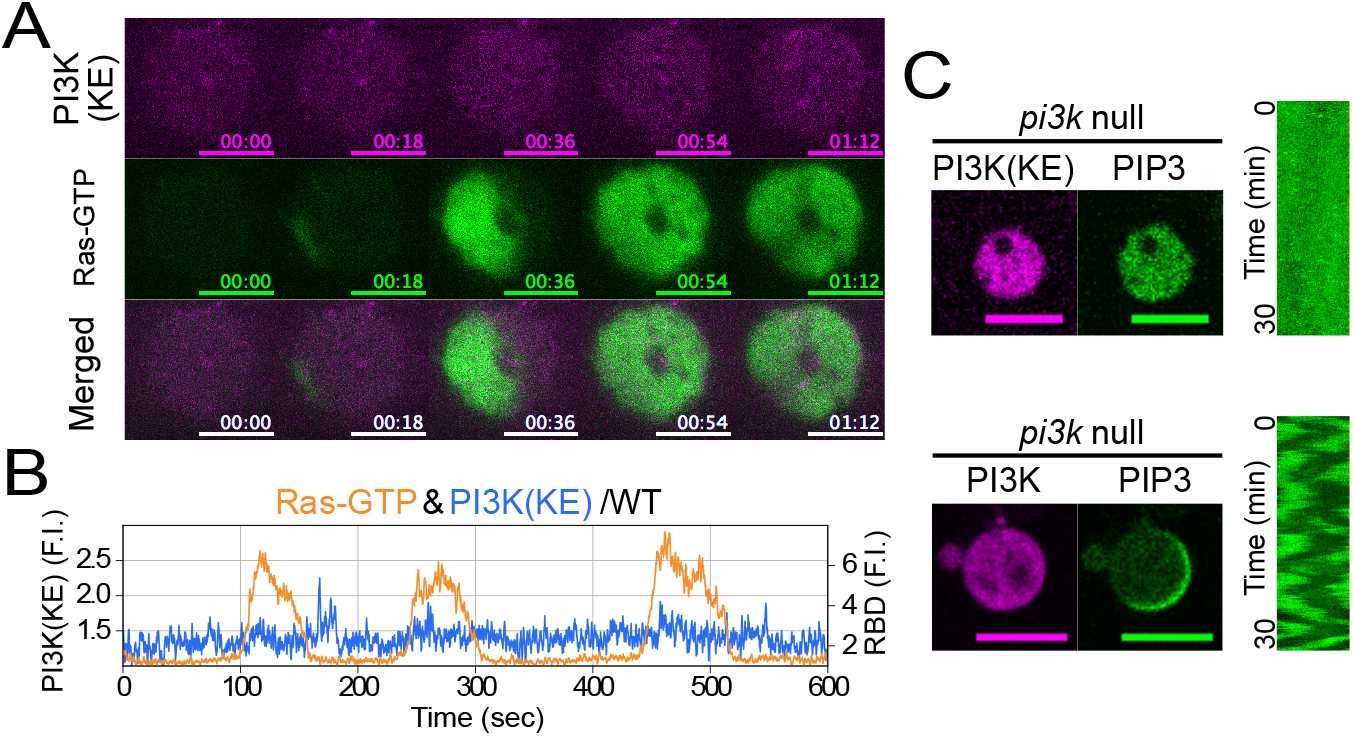
Ras triggers PIP3 waves by regulating the membrane translocation and activation of PI3K. (A) Simultaneous time-lapse imaging of RBD_Raf1_-GFP and PI3K2^K857,858E^-Halo-TMR taken by TIRFM. Scale bars, 5 *μ*m. Time format is “mm:ss”. (B) Typical examples of time trajectories of PI3K^K857,858E^ (blue) and Ras-GTP (orange). (C) Confocal images of the rescue experiments of *pi3k1-5* null strain with PI3K and PI3K^K857, 858E^. More than 100 cells were observed for each experiment. Scale bars, 10 μm.

### Feedback stabilizes Ras waves in the default state

In contrast to Ras waves being generated in *pi3k1-5* null strain (Fig 1), a previous study has shown that Ras waves were suppressed by the PI3K inhibitor LY294002 [13,37]. We investigated this apparent inconsistency between the genetic and pharmacological inhibition. When treated with 100 μM LY294002, the Ras-GTP waves vanished, as reported previously. However, with prolonged observation, we found that the Ras waves recovered gradually in the presence of LY294002 (Fig 4A and 4D, S8 Movie). The number of cells showing Ras waves transiently decreased to about 1% from about 84% (n=715), but finally recovered to about 36% (Fig 4B). Simultaneous observation of Ras and PIP3 revealed both waves were disappeared from most cells concomitantly, and that only the Ras waves were recovered by sixty minutes after treatment (Fig 4C). There were no remarkable effects by LY294002 on the Ras waves in *pi3k1-5* null cells (Fig 4B, S4A Fig). These results are consistent with our early conclusion that Ras wave generation is essentially independent of PIP3 production, additionally suggesting a novel mechanism that Ras, PI3K and PIP3 waves are tightly coupled in WT cells at the default state.

**Figure 4.**
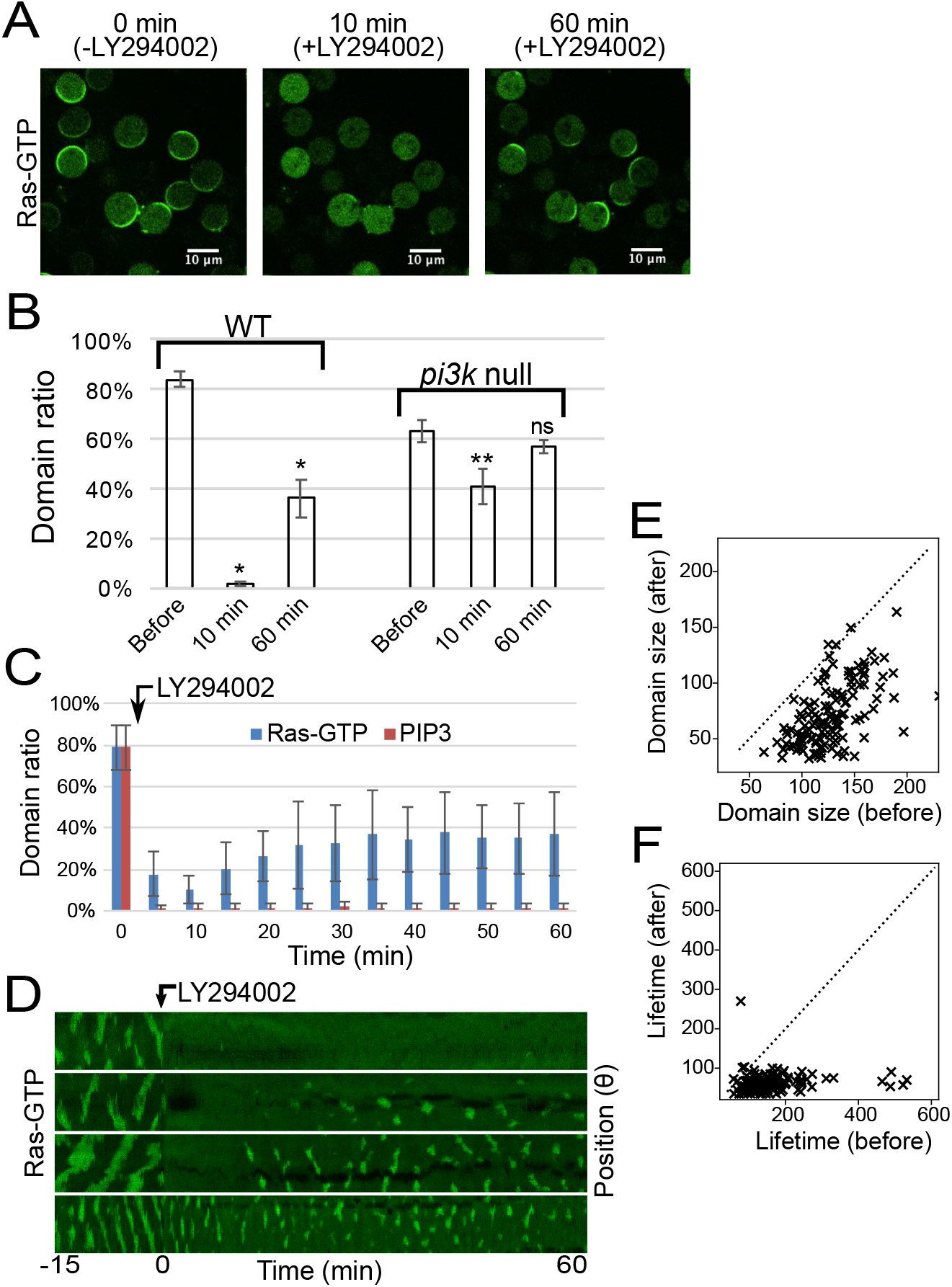
The effect of PI3K inhibition on Ras waves. (A) Response of Ras waves to a PI3K inhibitor (100 μM LY294002) before (left) and after 10 min (center) or 60 min (right) of treatment. (B) Ratio of the number of cells showing Ras-GTP domains before and after treatment with LY294002 in WT and *pi3k1-5* null strains. Data are mean ± s.d. of three independent experiments. More than 150 cells were counted in each experiment. (* P < 0.01, ** P < 0.01 Welch’s t-test; comparisons were made with “Before” column for the same cell type). (C) Ratio of the number of cells showing Ras-GTP or PIP3 domains after treatment with LY294002. Data are mean ± s.d. of four independent experiments. More than 40 PHD_AKT/PKB_-GFP and RBD_RAF1_-RFP doubly expressed cells were observed. (D) The kymographs show typical responses of Ras waves before and after LY294002 treatment. (E), (F) Distribution of the domain size or lifetime transition of the Ras wave pattern in each cell. Dotted lines indicate where Lifetime (before) equals Lifetime (after). These values were measured and averaged in each cell. 139 recovered cells were measured.

We noticed that the spatiotemporal patterns of Ras waves after LY294002 treatment were smaller and less continuous than those before the treatment (Fig 4D). Kymographs showed recovery of the Ras wave, although the Ras-GTP-enriched domains did not propagate continuously and instead underwent excitation transiently. The size and lifetime of the Ras-GTP-enriched domains became smaller and shorter with LY294002 treatment, respectively (Fig 4E and 4F). Similar defects in Ras waves were also observed in *pi3k1-5* null strain as described above (S1C Fig), but LY294002 treatment did not affect the size or lifetime of the domains in *pi3k1-5* null strain (S4C and S4D Fig). Thus, PI3K activity is not essential for the generation of activated Ras-GTP-enriched domains, but feedback from PIP3 to Ras enhances the excitability to stabilize the asymmetric signals.

### Interrelationship analysis of traveling waves in the Ras-PIP3 system

To reveal all intermolecular relationships relevant to Ras and PIP3 waves, we systematically analyzed the interrelationships between traveling waves of all possible combinations of two components among Ras-GTP, PI3K2, PTEN, PIP2 and PIP3. As a fluorescent probe for PIP2, GFP-tagged Nodulin (GFP-Nodulin) was used, which gave a brighter signal than the GFP-tagged PH domain derived from PLCδ1 (PHD_PLCδ1_-GFP) on the plasma membrane of *Dictyostelium* cells (S7 Fig, S7 Movie) [30,38]. We obtained phase diagrams from the average dynamics from the simultaneous TIRFM observations (Fig 5), which revealed all these components exhibited travelling waves along the membrane in a closely coupled manner (S5-S7 Fig, S6-S7 Movie). We found five characteristic features in their relationships. First, a phase diagram in Ras-GTP-PI3K2 coordinates showed that the amount of PI3K2 on the membrane was proportional to that of Ras-GTP and tight regulation on PI3K membrane recruitment by Ras-GTP (Fig 5A). The propotional relation explains the resemblance in the phase diagrams for Ras-GTP-PTEN and PI3K2-PTEN (Fig 5B-5E), for Ras-GTP-PIP3 and PI3K2-PIP3 (Fig 5C-5F), and for Ras-GTP-PIP2 and PI3K2-PIP2 (Fig 5D-5G). Second, the Ras-GTP-PTEN and PI3K2-PTEN diagrams exhibited a triangle shape with counterclockwise time progression (Fig 5B and 5E).

**Figure 5.**
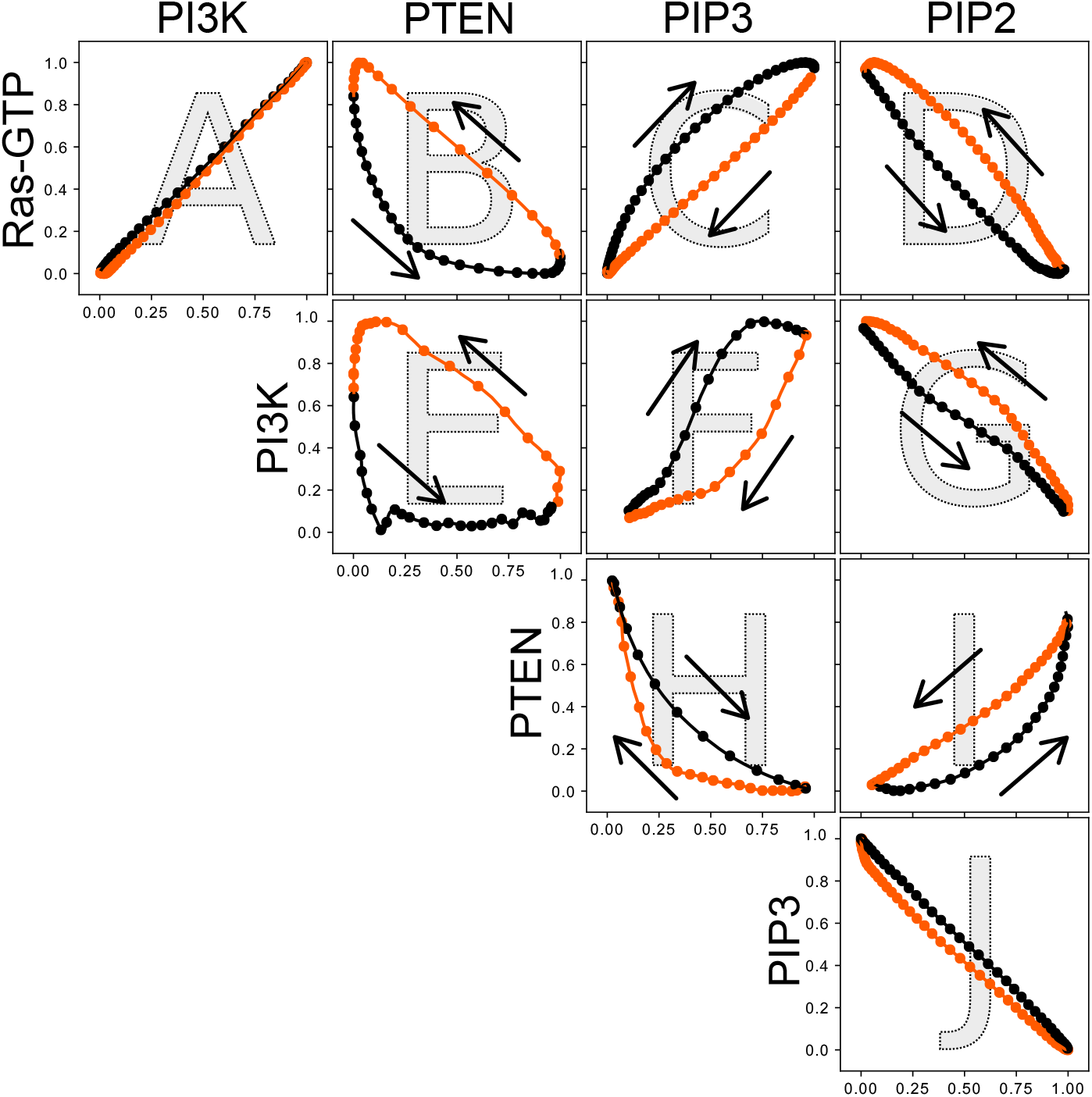
Interrelationship between traveling waves of Ras-GTP, PI3K, PTEN, PIP3 and PIP2. Average dynamics of two arbitrary components are shown as an orbit in the corresponding coordinates. Arrows indicate temporal progression. Black lines indicate increased phase along the horizontal axis and red lines indicate the decreased phase.

In the initial moment of the decreasing phase of the Ras-GTP/PI3K2 waves, the decrease in the amount of Ras-GTP and PI3K2 on the membrane was followed by a delayed increase in PTEN, revealing an order in the temporal changes of these molecules. Third, the Ras-GTP-PIP3 and PI3K2-PIP3 diagrams showed a characteristic ellipse-like trace with temporal clockwise progression, which is consistent with Ras/PI3K2 waves preceding PIP3 waves (Fig 5C and 5F). Fourth, the PIP3-PIP2 diagram exhibited an inverse relation with almost the same trace in the increasing and decreasing phases, suggesting that the total amount of PIP2 and PIP3 was almost constant during the wave propagation (Fig 5J, S7D Fig, S7 Movie). The inverse relation explains the mirror-image relations found between the Ras-GTP-PIP3 and Ras-GTP-PIP2 diagrams (Fig 5C and 5D), the PI3K2-PIP3 and PI3K2-PIP2 diagrams (Fig 5F and 5G), and the PTEN-PIP3 and PTEN-PIP2 diagrams (Fig 5H and 5I). Finally, the PIP3-PTEN and PIP2-PTEN diagrams exhibited a previously reported crescent shape trace with clockwise temporal progression, showing PTEN increase follows PIP3 decrease and PIP2 increase (Fig 5H and 5I) [13].

### Modeling and simulation of symmetry breaking in the Ras-PIP3 system

The dynamics of the components can be naturally explained and reconstituted in a numerical simulation based on a model described by a series of reaction-diffusion equations based on the experimental observations (Fig 6A, see *Methods* for detail). Ras excitability was modeled with positive and delayed negative feedbacks on Ras, two features commonly assumed in excitable systems and have been adopted in previous models for the self-organization of PIP3-enriched domains [3,13,29]. PI3K on the membrane is proportional to Ras-GTP levels and catalyzes PIP3 production according to a simple Michaelis-Menten (MM) type enzymatic reaction. PTEN is recruited to the membrane via interaction with PIP2 and excluded from the membrane by interaction with PIP3, phenomena that are also described by MM type binding reactions [13,39,40]. The observed feedback was modeled so that an assumptive regulator activates Ras in a PIP3-dependent manner (Fig 4). No other reaction pathways that cause PIP3 and PIP2 metabolism were taken into account in this simplified model because of the inverse relationship between PIP3 and PIP2.

**Figure 6.**
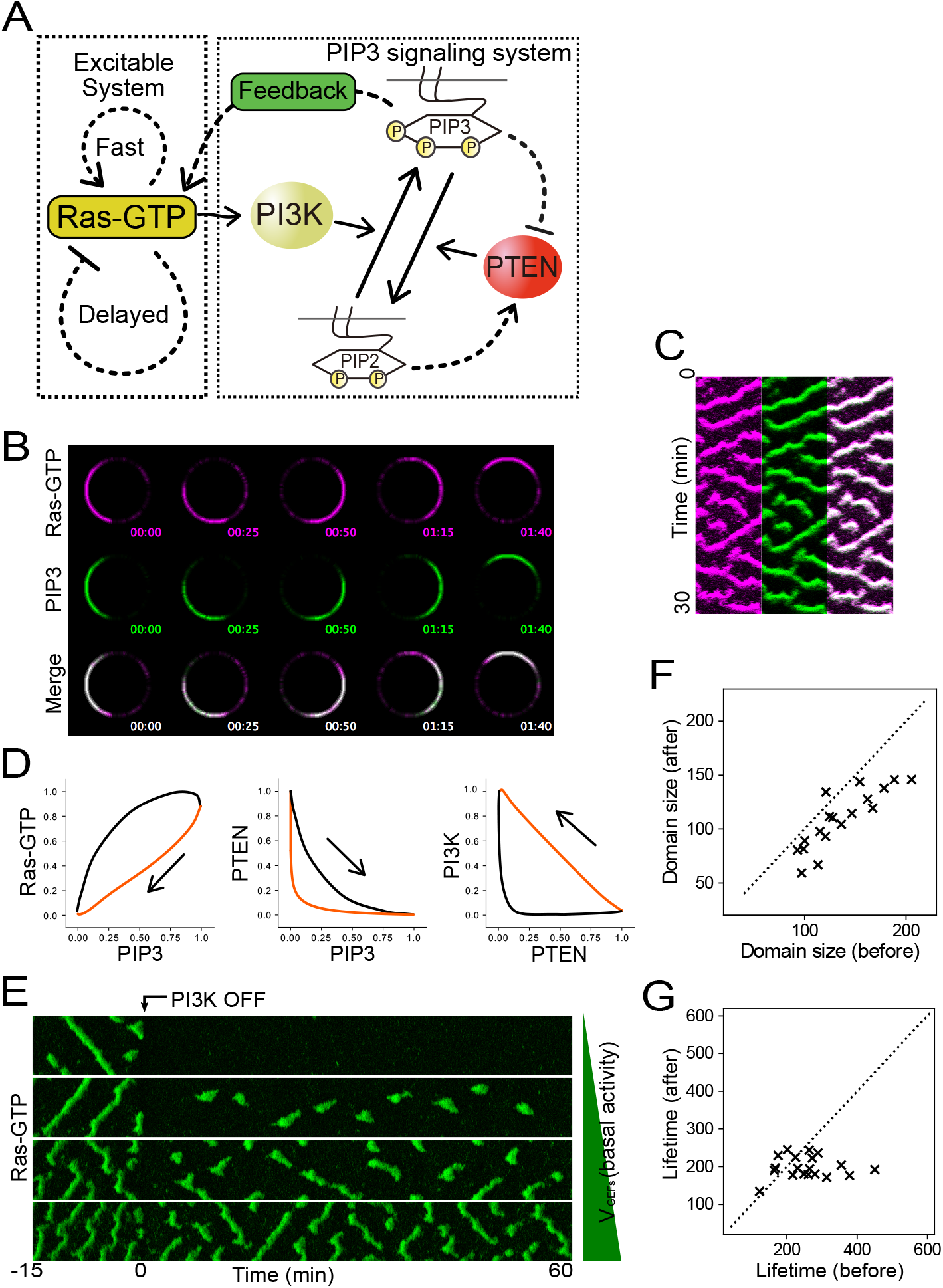
Spatiotemporal simulation of Ras/PIP3 wave formation. (A) Scheme of the Ras/PIP3 wave model used in this study. See text and *Methods* for details. (B) Ras-GTP and PIP3 traveling waves generated from the spatiotemporal stochastic simulation of the model shown in Fig 6A. *V_GEFs_* = 550 s^-1^. (C) Kymographs based on Fig 6B. (D) Phase diagrams generated from the deterministic simulation (see also S8 Fig). (E) Spatiotemporal stochastic simulation of the model and PI3K inhibition. Kymographs show waves before and after PI3K inhibition. After PI3K inhibition, Ras waves vanished immediately, but recovered after a few minutes depending on the basal activity (*V_GEFs_* = 400, 420, 450, 500 s^-1^). (F), (G) Distribution of the domain size and lifetime transition of the Ras wave pattern in each cell. Dotted lines indicate where Lifetime (before) equals Lifetime (after). (*V_GEFs_* = 400 ~ 600 s^-1^ in 10 increments).

We performed the simulation and confirmed the model can reconstruct all the traveling waves in a manner consistent with the experimental observations (Fig 6B and 6C). By assuming no PI3K activity, the one-dimensional spatiotemporal simulation reconstructed Ras-GTP traveling waves without PIP3 waves due to the excitability (S8A Fig). By applying PI3K activity, the traveling waves of PIP3 also appeared in a co-localized manner with Ras-GTP (Fig 6B and 6C). Under this situation, the temporal oscillations of Ras-GTP, PI3K, PTEN, PIP2 and PIP3 were all reconstructed, and phase diagrams plotted in the corresponding coordinates were consistent with the experimental observations showing the five characteristic features in their relationships (Fig 6D, S8B Fig). Thus, the interrelated oscillations of the Ras-GTP-PIP3 system can be explained by our model, in which activated Ras triggers the other waves through its interaction with PI3K.

We next examined the effects of positive feedback from PIP3 on Ras wave generation by turning on/off PI3K activity under broader conditions of the simulation. We performed the simulation by changing *V_GEFs_*. In the presence of PI3K activity, i.e. positive feedback from PIP3 to Ras, travelling waves were generated across a wide range of *V_GEFs_* (Fig 6E). Upon the inhibition of PI3K and thus no feedback regulation, Ras wave dynamics became dependent on *V_GEFs_.* Under intermediate conditions, Ras waves exhibited a recovery after a transient loss. The transient loss and recovery can be explained as follows. In our model, larger *V_GEFs_* causes higher Ras-GTP basal levels and thereby a smaller distance to the threshold in the excitable system. At intermediate *V_GEFs_,* PI3K inhibition puts the system below the threshold, leading to a loss of Ras excitation. Then, the delayed negative feedback working on Ras gradually loses its suppressive activity, allowing the system to exceed the threshold and recover Ras excitation. During the recovery process, molecular noise helps stochastic crossing of the threshold, since the recovery was not reconstructed in deterministic simulations (S8C Fig). The Ras excitation recovery can be also reproduced by increasing *V_GEFs_* with a lag time after PI3K inhibition, an effect that can be regarded as changing the expression levels of the components responsible for Ras excitation. Furthermore, we performed kymograph analysis of the Ras-GTP dynamics obtained from the simulation and found that the size and lifetime of the Ras-GTP domain became smaller and shorter after the inhibition of PI3K activity, respectively, which is in good agreement with the experimental observations (Fig 6F and 6G). These results suggest that feedback regulation from downstream pathways contribute to the maintenance of Ras excitability.

## Discussion

Live-cell imaging analysis of asymmetric signal generation in Ras and the PIP3 signaling pathway demonstrates that there exist at least three characteristics of the self-organization process. First, Ras GTPase has a central role in the asymmetric signal generation to initiate traveling waves independently of downstream pathways including the PIP3 signaling pathway (Fig 1). Second, Ras waves trigger the self-organization of the traveling waves of all components observed in the PIP3 signaling pathway through PI3K activation (Fig 2, 3 and 5). Third, feedback on Ras GTPase from the downstream molecules stabilizes the asymmetric signal generation (Fig 4). Overall, these experimental observations demonstrate that Ras GTPase has excitable dynamics and governs spatiotemporal dynamics in the phosphatidylinositol lipids signaling pathway for asymmetric signal generation.

In the context of chemotaxis, the excitability of Ras GTPase provides an amplification of gradient signals detected by chemoattractant receptors. The molecular mechanisms for chemotaxis are evolutionally conserved among various cell types including mammalian neutrophils and social amoeba *Dictyostelium discoideum,* in which spontaneous cell motility is biased directionally along chemical gradients [5,30]. Gradient sensing has been explained theoretically by a local excitation and global inhibition (LEGI) model combined with a signal transduction excitable network (STEN) model [9]. LEGI can detect gradient signals, and the output is generated at the membrane that faces the higher chemoattractant concentration. STEN works as an amplifier of LEGI output to generate a localized signal in an all-or-none fashion [41]. Our findings are consistent with the idea that Ras GTPase can be regarded as an essential part of the STEN interface with LEGI. Spatial differences in Ras activity derived from chemoattractant receptors can be amplified in an all-or-none manner along chemical gradients. That is, the internal signals that are generated spontaneously are integrated with external signals derived from chemoattractant receptors through the spatiotemporal regulation of Ras GTPase, leading to biased asymmetric signals along the chemical gradients for chemotaxis.

Ras GTPases and the regulatory network make up a complex system comprised of various Ras GTPase family proteins, RasGEFs and GTPase activating proteins (GAPs). In *Dictyostelium discoideum* cells, at least 14 Ras GTPase family proteins, 25 GEFs, and 14 GAPs are estimated from the genome sequence [6,42]. Our findings here demonstrate that the Ras GTPase regulatory network has excitable dynamics, meaning that it has a threshold for excitation. In general, GEFs and GAPs regulate Ras GTPase positively and negatively, respectively. That is, the inputs from GEFs and GAPs can be regarded as excitatory and inhibitory signals for Ras GTPase. These signals are integrated through the regulation of Ras GTPase activity and cause excitation when the network exceeds the threshold. This concept is analogous to a neuron in neuronal networks, because the neuron has characteristics of an excitable system in which excitatory and inhibitory signals derived from presynaptic cells are integrated in the postsynaptic cell during the signal processing. It should be noted, however, that the hierarchical level of Ras GTPases differs from that of neurons. From the viewpoint of molecular signal processing in excitable systems, it is important to clarify the circuit structure of the Ras GTPase regulatory network in order to provide mechanistic insights into cellular decision-making processes. Because Ras and the phosphatidylinositol lipids signaling pathway are involved in oncogenesis and metastasis, understanding the mechanisms that regulate the threshold of the Ras excitable system are important issues in the biological and medical sciences.

## Materials and Methods

### Cell culture and constructs

*Dictyostelium Discoideum* wild-type Ax2 was used as the parental strain except when *gc* null strain (Ax3) was used. Cells were transformed with plasmids based on the vector series pDM or pHK12 by electroporation. Plasmids were generated by Ligation or In-Fusion (TOYOBO, TaKaRa). All constructs were sequenced before transformation. To visualize the localization of PI3K2 and its mutants, Halo-tag^®^ fused PI3K2 was expressed. A Ras binding mutant of PI3K2 (PI3K2^K857E, K858E^) and truncations of PI3K2 N-terminal (PI3K2^Δ561-1859^) were obtained by PCR (TOYOBO). To observe the spatiotemporal dynamics of phosphatidylinositol lipids and their related enzymes, pairs of PHD_AKT/PKB_, Nodulin, PTEN, PI3K and RBD_Raf1_ tagged with GFP, RFP or Halo-TMR were co-expressed. Cells were cultured axenically in HL-5 medium containing G418 (20 mg/ml), Blasticydin S (10 mg/ml) or hygromycin (50 mg/ml) at 21 °C [43].

### Preparation for live cell imaging

For imaging experiments, cells were washed in 1 ml development buffer without Ca^2+^ or Mg^2+^ (DB-; 5 mM Na_2_HPO_4_ and 5 mM KH_2_PO_4_) twice and starved in 1 ml development buffer (DB; DB-, 2 mM MgSO_4_ and 0.2 mM CaCl_2_) for 3 to 4 hours at a density of 5×10^6^ cells per ml on a 35 mm dish. Halo-tag^®^ (Promega)-fused proteins were stained with 2 *μ*M Halo-tag^®^ TMR ligand for 30 min and washed 5 times with 1 ml DB-before observation.

### Imaging with confocal microscopy

Confocal imaging was performed using an inverted microscope (ECLIPSE Ti; Nikon) equipped with a confocal unit (CSU-W1; Yokogawa). Laser sources for 488 nm and 561 nm excitation light were solid-state CW lasers (OBIS 488NM X 50MW and OBIS 561NM X 50MW, respectively; COHERENT). Time-lapse images were acquired through a 60 × oil immersion objective lens (CFI Apo TIRF 60X Oil, N.A. 1.49; Nikon) with an EM-CCD camera (iXon3 897; Andor). Cells were transferred to a 35 mm Glass Base Dish (Grass 12 φ, 0.15-0.18 thick; IWAKI) and suspended in 200 *μ*l DB with 4 mM caffeine and 5 *μ*M Latrunculin A (SIGMA). Inhibitors were added after 15 min treatment with caffeine and Latrunculin A. Latrunculin A (2 mM), LY294002 (40 mM), Torin2 (1 mM) and BPB (10 mM) in DMSO were diluted to the final concentration (DMSO ≤ 1%). Time-lapse images were obtained at 200 ms exposure for each channel at 2 s intervals (488 nm laser power was ~50 *μ*W, and 561 nm laser power was ~150 *μ*W).

### Imaging with TIRFM

TIRF imaging was performed using an inverted microscope equipped with a handmaid prism-less TIR system [44]. Laser sources for 488 nm and 561 nm excitation light were solid-state CW lasers (SAPPHIRE 488-20 and Compass 561-20, respectively; COHERENT). Lasers were guided to the back focal plane of the objective lens (CFI Apo TIRF 60X Oil, N.A. 1.49; Nikon) through a back port of the microscope. TIR and EPI illumination were switched by tilting the incident angle of the lasers. The separated images were passed through dual band laser split filter sets (Di01-R488/561-25×36, Di02R561-25×36, FF01-525/45-25 and FF01-609/54-25; Semrock) and captured by two EM-CCD cameras (iXon3 897; Andor) equipped with 4 × intermediate magnification lenses (VM Lens C-4 ×; Nikon). Cells were transferred to a cover glass (25 mm radius, 0.12-0.17 thick; MATSUNAMI) that was fixed on a chamber (Attofluor^®^ Cell Chamber; Molecular Probes). The cover glass was washed by sonication in 0.1 M KOH for 30 min and in 100% EtOH for 30 min twice in advance. Cells were treated with 4 mM caffeine and 5 *μ*M Latrunculin A in DB for 15 min. Time-lapse images were obtained at 100 ms exposure for 10 min (488 nm laser power was ~20 *μ*W, and 561 nm laser power was ~20 *μ*W). Time-lapse images were smoothed with a 1 s time window to reduce shot noise.

### Image processing and analysis

The adjustment of images from a dual camera was performed on software by using objective micrometer images as a standard. The average error of the position after adjustment was less than 66 nm^2^. Leakage from one channel to the other was calibrated as:

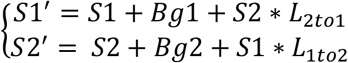

*S*_1_ and *S*_2_ indicate the fluorescence signals from the two channels. *S*’ indicates the detected signal, *Bg* indicates the background noise and *L* indicates the ratio of leakage from one signal to the other channel. Leakage parameters for GFP, RFP and TMR were obtained in advance.

The spatiotemporal dynamics were analyzed from the time trajectories of the membrane localization. For TIRF time-lapse images, time trajectories were the time-axis profiles of the average intensity of the ROI. We defined the ROI as a circle with 0.2 *μ*m (3 pixels) radius. In the case of confocal time-lapse images, the fluorescence intensities were analyzed as previously described [13]. Fluorescence intensity along the rounded cell contours were measured in each frame. The intensity profiles were plotted against the angle θ and time as a 2D pattern. The line profile along the time axis was defined as the time trajectory of the membrane localization. This method can reduce the effect of stage drift.

### Time trajectory analysis

Raw trajectories were extracted from TIRF images as described above. Auto- and cross-correlation functions were calculated from the trajectories. To obtain the average trajectories shown in Fig 2 and 3, we calculated the local maximums and minimums of the first derivative of the raw trajectories at the centers of the increasing and decreasing phases, respectively. Next, we extracted short trajectories of 60 s before and after the calculated centers from the raw trajectories. Finally, we aligned the short trajectories at their centers, normalized them by z-scores and calculated the average trajectories. The phase diagrams were drawn from the average trajectories.

### Domain size and duration time analysis

The domain size and duration time were calculated from binarized kymographs. Kymographs were smoothed by a mean filter with a 9-pixel window and binarized to separate the domain and background. Then we detected each domain and defined the duration time as the size of the domain along the time axis and the domain size as the average spatial size of the domain in each time step. We ignored domains smaller than 32° or shorter than 30 s.

### Reaction diffusion model and numerical simulations

To reconstruct the spatiotemporal dynamics of the Ras/PIP3 wave patterns, we constructed a combined model of the Ras and PIP3 signaling systems (Fig 6A). As described in the main text, excitability and pattern formation derive from Ras and its regulatory network. The PIP3 signaling system follows Ras-GTP, and feedback from PIP3 to Ras maintains Ras excitability.

First, we constructed a model of the PIP3 signaling system as described below.

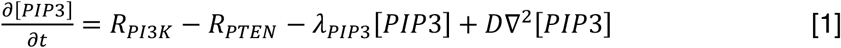

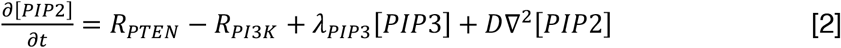

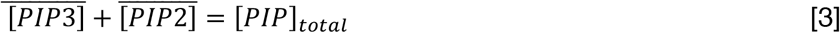

PIP3 is generated from PIP2 phosphorylation by PI3K (*R_PI3K_*) and dephosphorylated into PIP2 by PTEN (*R_PTEN_*) on the membrane [1], [2]. The PTEN-independent PIP3 degradation rate, *λ_PIP3_*, is also introduced because the degradation of PIP3 is observed even in *pten* null strain [27]. The sum of PIP3 and PIP2 concentrations was made constant [3] (Fig 5J). PIP3 and PIP2 diffuse on the membrane independently with the diffusion term *D*∀^2^.

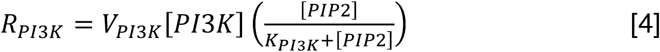

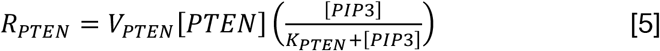

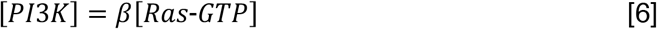

The PIP2 phosphorylation reaction (*R_PI3K_*) and PIP3 dephosphorylation reaction (*R_PTEN_*) were described as Michaelis-Menten type enzymatic reactions, in which each term is composed with the maximum reaction rate (*V_PI3K_*, *V_PTEN_*) and Michaelis constant (*K_PI3K_*, *K_PTEN_*) [4], [5]. Because PI3K shuttles between the cytosol and plasma membrane dependently on Ras-GTP interactions, we introduced Ras-GTP-dependent PI3K membrane translocation and Ras-GTP-dependent PI3K activation [6]. *β* indicates the membrane association rate of PI3K, because the membrane translocation of PI3K is proportional to the Ras-GTP level (Fig 5A).

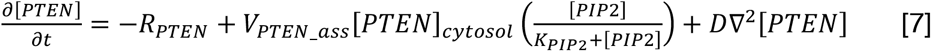

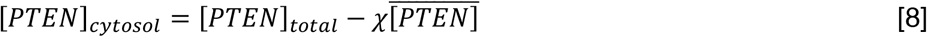

PTEN also shuttles between the cytosol and plasma membrane and shows a mutually exclusive localization pattern with PIP3 (Fig 5H). To satisfy these constraints, we introduced PIP3-dependent exclusion of PTEN from the membrane [13,14,39]. We assumed interaction between PTEN and PIP3 results in PTEN dissociation from the membrane and the dissociation rate has the same rate with PIP3 dephosphorylation, *R_PTEN_* [7]. Additionally, we introduced PTEN recruitment to the membrane by interaction with PIP2 [39] as dependent on the maximum reaction rate, *V_PTEN_ass_*, and Michaelis constant, *K*_*PIP*2_ [7]. Unlike the case of PI3K, the cytosolic concentration of PTEN drastically changes before and after membrane translocation [27]. Therefore, we considered the cytosolic concentration of PTEN. As described previously, we assumed the cytosolic PTEN concentration is uniform inside the cytosol and described by equation [8] [13].

Second, we constructed the Ras signaling regulation model as an excitable network. As a template for the excitable network model, we applied the theoretical model of self-organization on the membrane reported previously [14], which can reconstruct well the excitable responses of a Ras/PIP3 signaling system with a minimum number of elements.

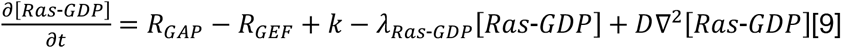

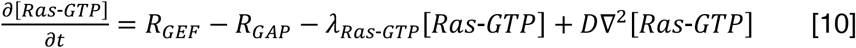

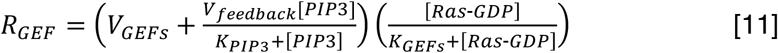

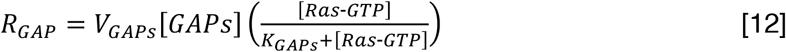

In this model, we introduced a positive regulator and negative regulator of Ras as GEFs and GAPs, respectively. Ras is activated by reaction *R_GEFs_* and inactivated by reaction *R_GAPs_*. Additionally, Ras-GDP is supplied and degraded at the rates *k* and *λ_RasGDP_*, respectively [9]. Ras-GTP is degraded at *λ_RasGTP_* and supplied only by the GTP exchange reaction of Ras-GDP [10]. *R_GEFs_* is described as a Michaelis-Menten type enzymatic reaction composed of two terms [11]. One term defines the basal activity of Ras and the other defines feedback from PIP3. Each term is composed of the maximum reaction rate (*V_GEFs_, V_Feedback_*[*PIP3*]/(*K_PIP3_* + [*PIP3*])) and Michaelis constant *K_GEFs_*. This formulation is based on the experimental observation that PIP3 production influences the formation of Ras waves (Fig 4). *R_GAPs_* is described as an enzymatic reaction with maximum reaction rate *V_GAPs_* and Michaelis constant *K_GAPS_* [12].

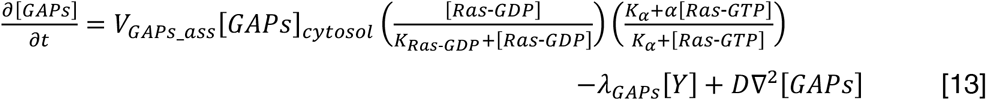

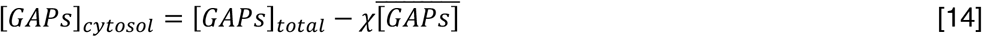

In order to promote the formation of a Ras-GTP localization patch, we introduced positive regulation from Ras-GDP and negative regulation from Ras-GTP on the recruitment of GAPs to the membrane. These regulations are composed of two positive-feedback loops and result in a mutually exclusive relationship between Ras-GTP and GAPs, which stabilizes spatially restricted Ras-GTP localization patches. Positive regulation from Ras-GDP is described as a Michaelis-Menten type enzymatic reaction composed of a maximum reaction rate (*V_GAPs_ass_*) and Michaelis constant (*K_Ras-GDP_*), and negative regulation is described as (*K_α_* + *α*[*Ras-GTP*])/(*K_α_* + [*Ras-GTP*]) following a previous study [13]. Additionally, we introduced the degradation of GAPs at rate *λ_GAPs_* and the cytosolic concentration as described in equation [13].

Detailed methods for the numerical simulations are described previously [14]. We evaluated a one-dimensional system with 100 grids along the membrane to reconstruct the results of the kymographs. The radius of the cells was chosen to be 5 *μ*m with a constant time step Δ*t* = 0.005. For the stochastic simulation, we used the τ-leap method. The spatiotemporal dynamics are described by the reaction diffusion equation described above.

The parameters are summarized in S1 and S2 Table. First, we set the parameters shown in S1 Table, which are related to the PIP3 signaling pathway. We performed a simulation in which the concentration term [Ras-GTP] was fixed to the value from the experiments and adjusted the parameters to satisfy the relationships between PIP3, PI3K and PTEN to the experimental results shown in Fig 5. Next, we set the parameters related to the Ras wave pattern formation in the condition without feedback from PIP3 by a simulation in which the feedback term *V_feedback_* was fixed to zero. Finally, we modulated *V_feedback_* and *V_GEFs_* to satisfy the experimental results.

Kymographs were generated from the simulations based on this model and parameters. The parameter value of PI3K activity, V_Pl3k_, was modulated during the simulation. Then, we set the value of V_pl3K_ to 0 s^-1^ (PI3K off) or 12 s^-1^ (PI3K ON) in the PI3K on/off simulation shown in Fig 6E. The value of Ras basal activity, *V_GEFs_*, was also changed in each simulation. We varied the values of *V_GEFs_* from 400 to 600 s^-1^ in 10 increments and analyzed all kymographs to obtain distributions of the domain size and duration time.

## Data availability

Data supporting the findings of this study are available from the corresponding authors upon request.

## ACKOWLEDGEMENTS

GFP-Nodulin was kindly provided by Y. Miao and P. N. Devreotes (Johns Hopkins University School of Medicine, Baltimore). We thank G. Gerisch (Max Planck Institute for Biochemistry, Germany) for the LimEΔcoil-RFP construct, R. A. Firtel (University of California, San Diego) for the RBD-GFP construct, T. Uyeda (National Institute of Advanced Industrial Science and Technology, Ibaraki, Japan) for the PH_AKT/PKB_-GFP construct, R. R. Kay (MRC Laboratory of Molecular Biology, Cambridge, UK) for the *pi3k1-5* null cells, NBRP for the *rasC* null cells and *rasG* null cells, P. J. Van Haastert (Department of Cell Biochemistry, University of Groningen, Netherlands) for the *gc* null cells, M. Okada (Laboratory of Cell Systems, Osaka University, Japan), T. Shibata (Riken BDR, Japan) and members of the Ueda labs for helpful suggestions. This work was supported by AMED-CREST (JP17gm0910001) from Japan Agency for Medical Research and Development, AMED and by Grant-in-Aid for JSPS Research Fellow (17J02075) from Japan Society for the Promotion of Science, JSPS.

## Supporting Information

**S1 Fig.**
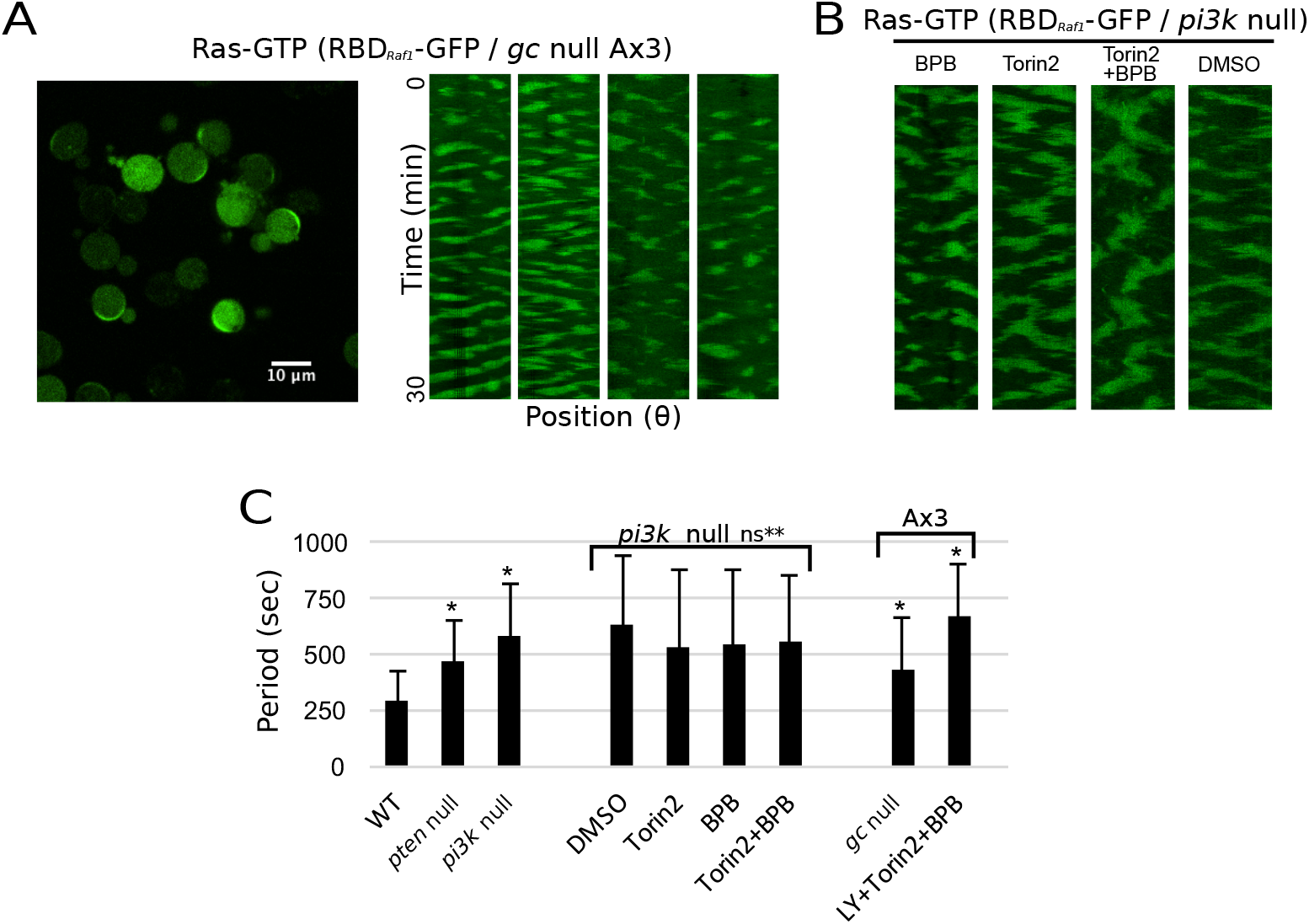
Ras waves in mutants. (A) Confocal images of Ras waves in AX3 background *gc* null cells expressing RBD_Raf1_-GFP. Kymographs shows typical Ras wave patterns. (B) Kymographs of *pi3k1-5* null cells treated with 2 *μ*M BPB, 10 *μ*M Torin2, a combination of them and 1% DMSO as a control. (C) Ras wave periods were calculated from the auto-correlation function of more than 20 cells and averaged. (* P < 0.01 Welch’s t-test against WT; ns** P > 0.01 Welch’s t-test against *pi3k1-5* null).

**S2 Fig.**
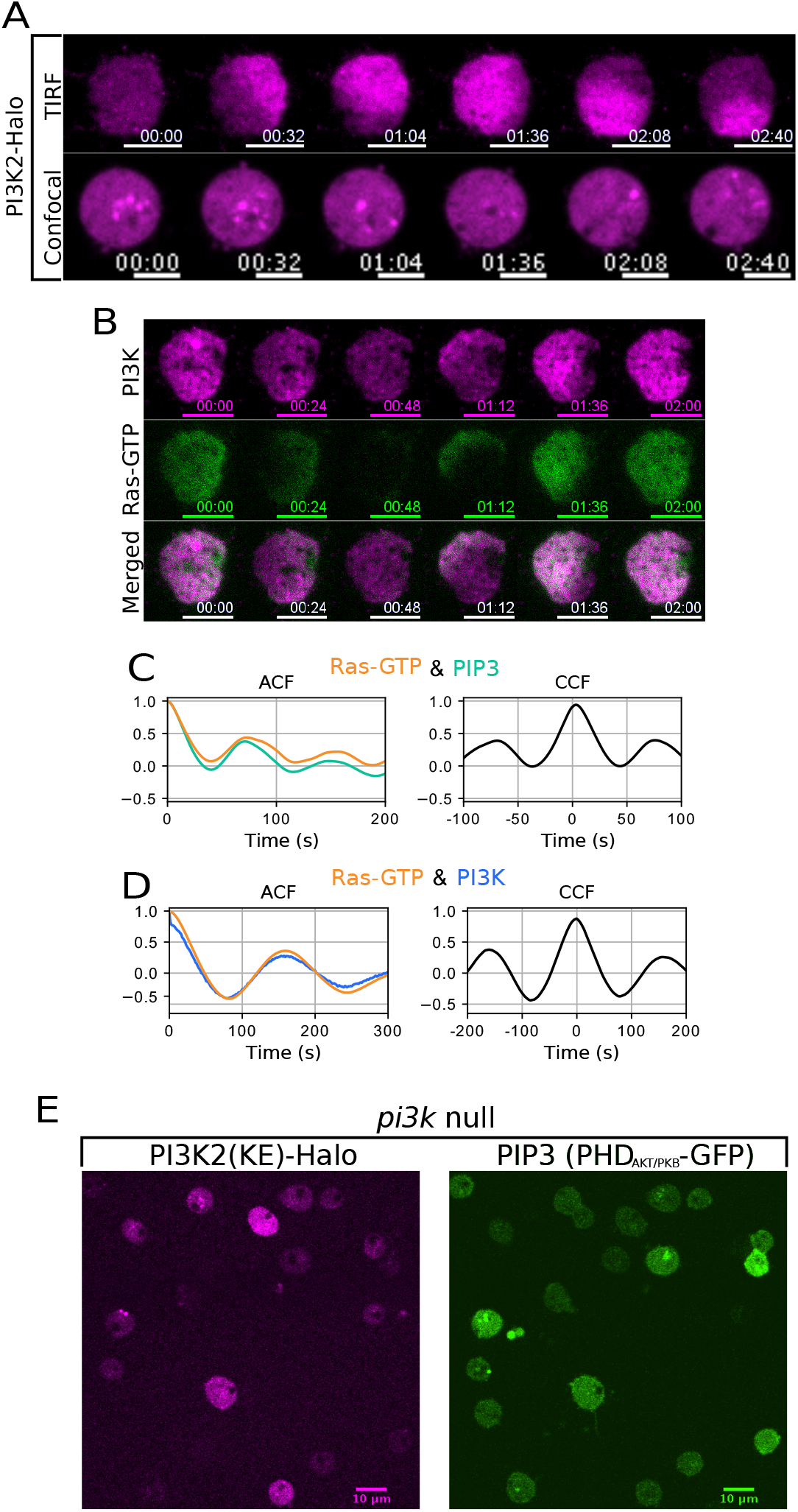
TIRF and confocal imaging of PI3K. (A) Time-lapse imaging of PI3K (PI3K2-Halo-TMR) taken by a TIRF microscope (top) and confocal microscope (bottom). Scale bars, 5 *μ*m. (B) Simultaneous time-lapse imaging of RBD_Raf1_-GFP and PI3K2-Halo-TMR taken by TIRFM. Scale bars, 5 *μ*m. Time format is “mm:ss”. (C), (D) Auto- and cross-correlation functions of the trajectories shown in Fig 2C and 2E, respectively. (E) Confocal images of *pi3k1-5* null strain expressing PI3K(KE)-Halo and PHD_AKT/PKB_-GFP. PIP3 domain formation is not rescued in these cells.

**S3 Fig.**
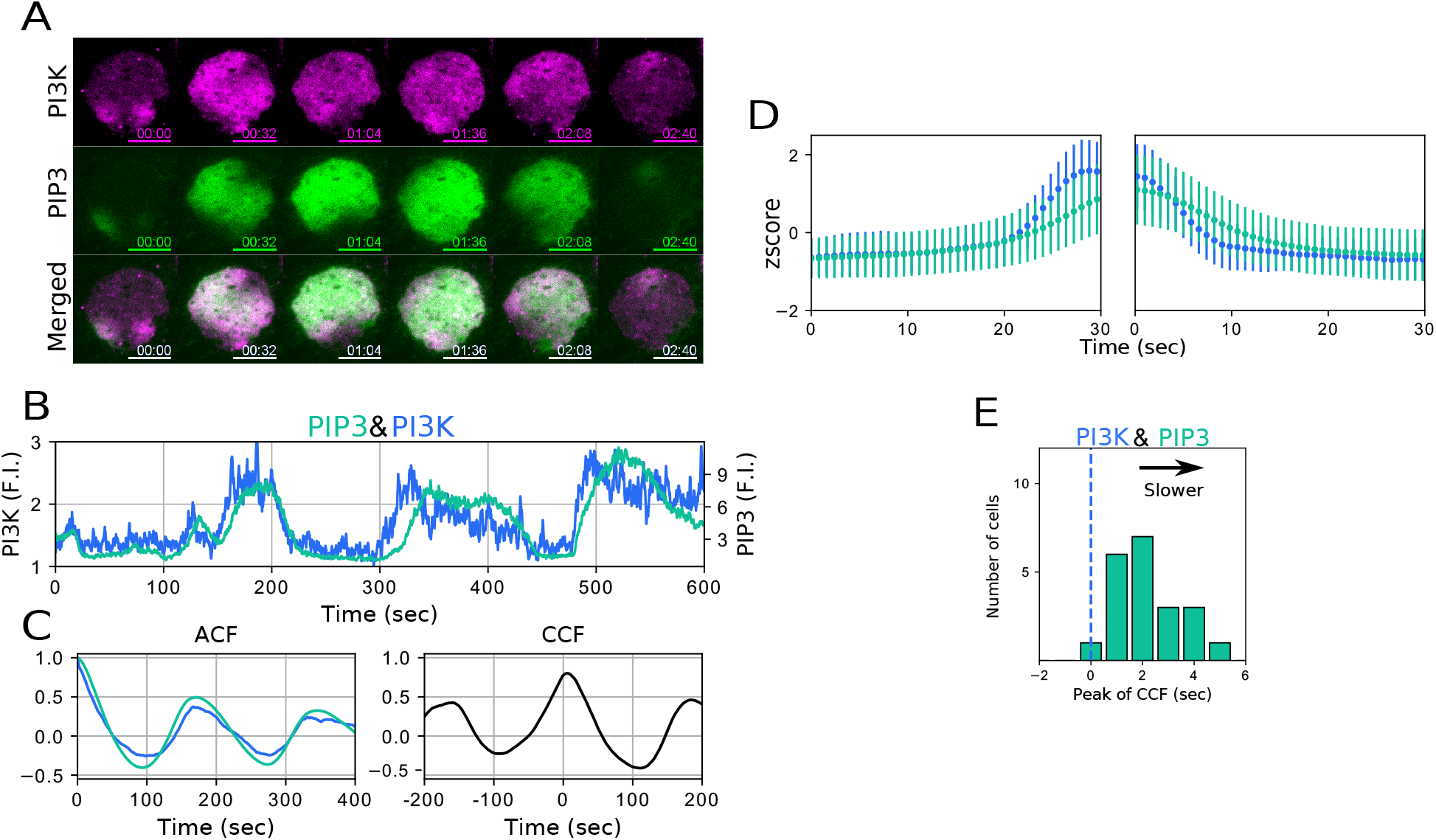
Simultaneous observation of PI3K and PIP3. (A) Simultaneous time-lapse TIRFM imaging of PI3K2-Halo-TMR and PHD_AKT/PKB_-GFP. (B) Typical examples of time trajectories of PI3K (blue) and PIP3 (green). (C) Auto- and cross-correlation functions of the trajectories shown in B. (D) Average trajectories of the increase phase (left) and decrease phase (right). Data are the mean ± s.d. from more than 20 cells. (E) Distribution of peak times of the crosscorrelation functions. Dotted lines indicate time zero. The average peak value is 2.3 ± 1.1 s.

**S4 Fig.**
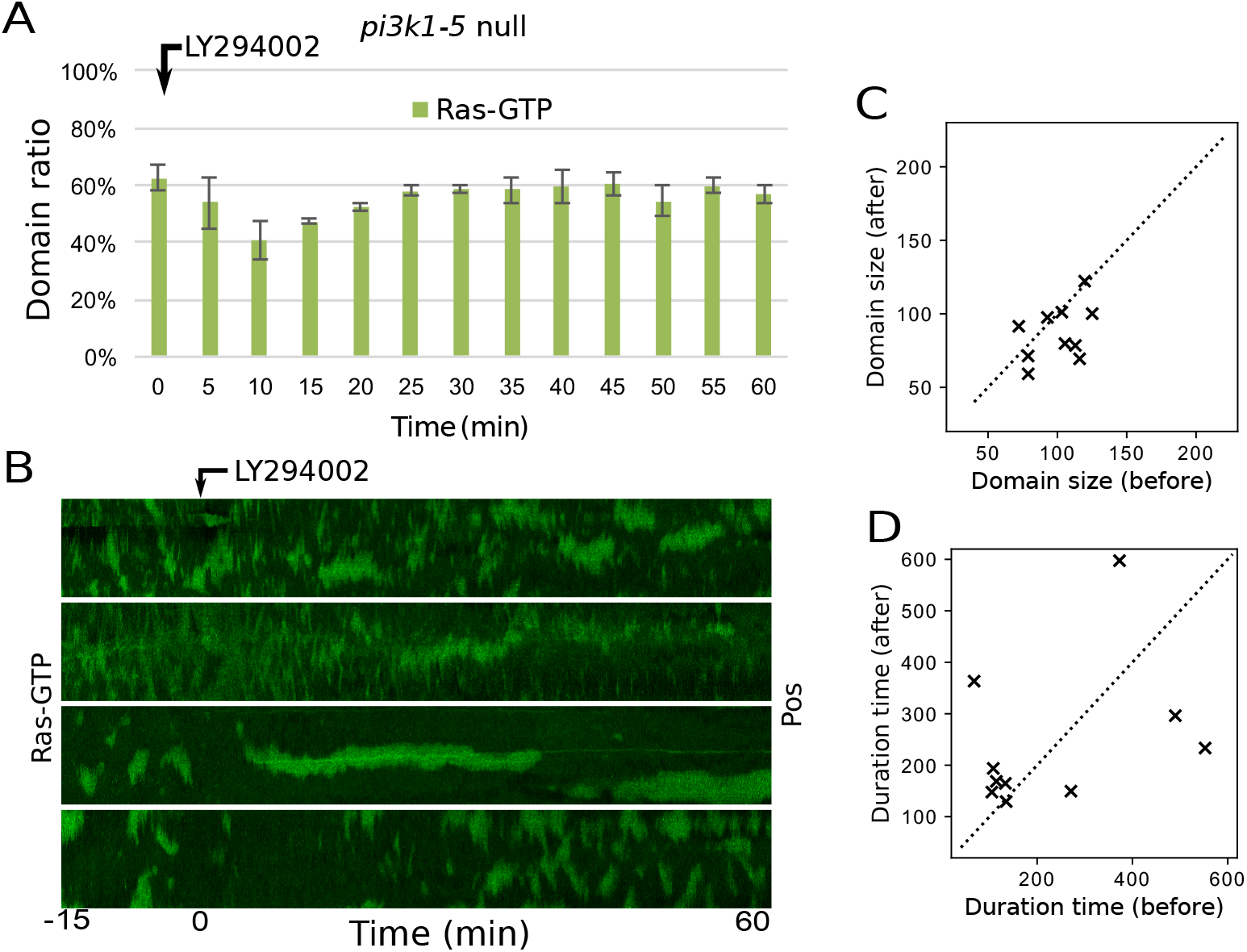
Ras waves with LY294002 treatment in *pi3k1-5* null cells. (A) Ratio of the number of cells showing Ras-GTP domains after treatment with LY294002 in *pi3k1-5* null cells. (B) The kymograph shows a typical response of a Ras wave in *pi3k1-5* null cells before and after treatment with LY294002. (C) Distribution of the domain size of the Ras wave pattern in each cell. 12 cells were measured. (D) Distribution of the duration time of Ras wave patterns in each cell.

**S5 Fig.**
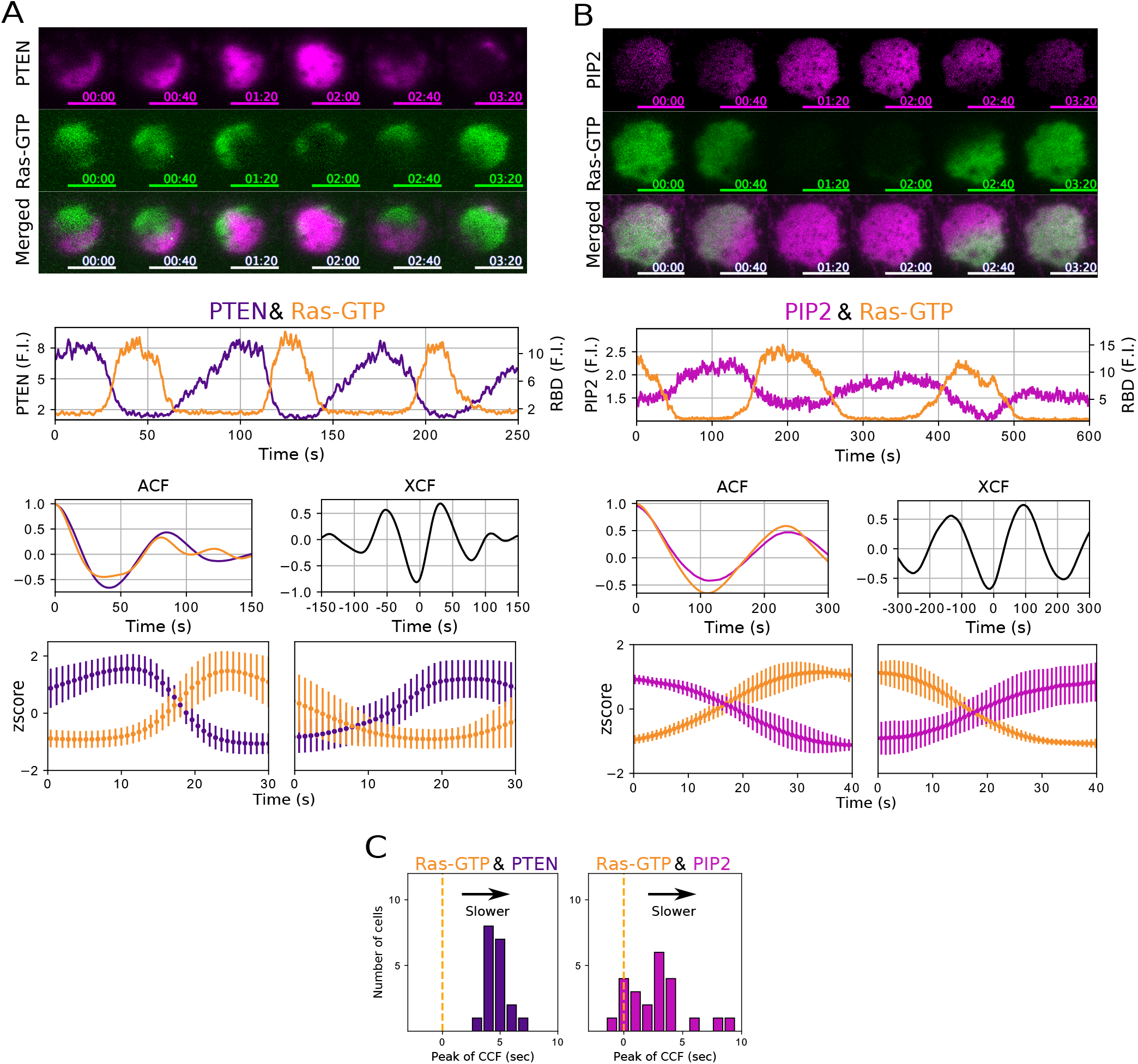
Simultaneous observation of Ras-GTP and PTEN or PIP2. Simultaneous time-lapse TIRFM imaging, typical time trajectories, and auto- and cross-correlation functions of (A) PTEN-Halo-TMR and RBD_Raf1_-GFP and (B) RFP-Nodulin and RBD_Raf1_-GFP. (C) Distribution of peak times of the cross-correlation functions. Dotted lines indicate time zero. The average peak value of Ras-GTP against PTEN is 4.7 ± 0.8 s and Ras-GTP against PIP2 is 2.7 ± 2.4 s.

**S6 Fig.**
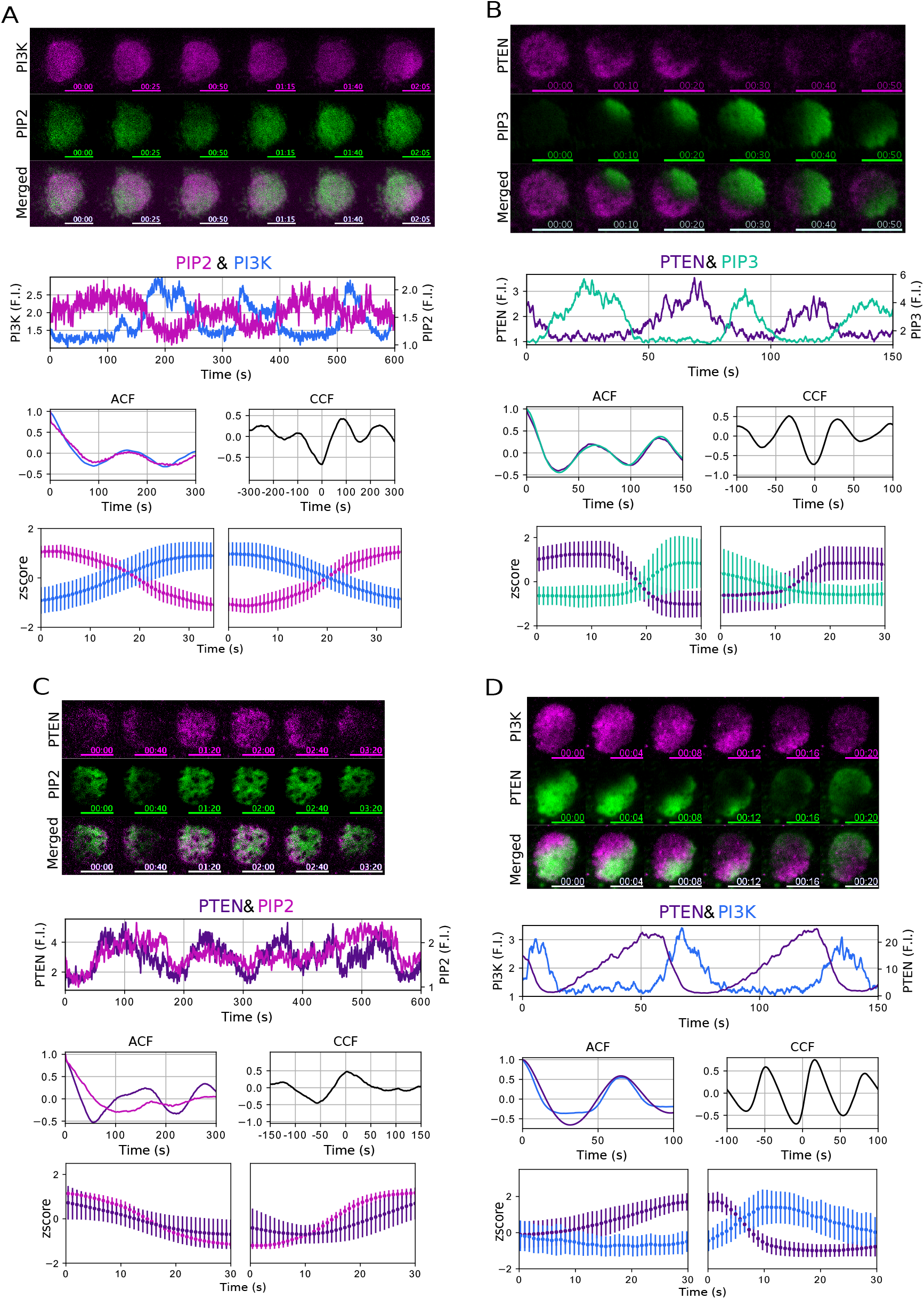
Temporal relationship between PI3K, PTEN, PIP3 and PIP2. Simultaneous time-lapse TIRFM imaging, typical time trajectories, auto- and cross-correlation functions and average trajectories of PTEN-Halo-TMR and RBDRaf1-GFP (A) PI3K2-Halo-TMR and GFP-Nodulin, (B) PTEN-Halo-TMR and PHD_AKT/PKB_-GFP, (C) PTEN-Halo-TMR and GFP-Nodulin and D, PI3K2-Halo-TMR and PTEN-GFP.

**S7 Fig.**
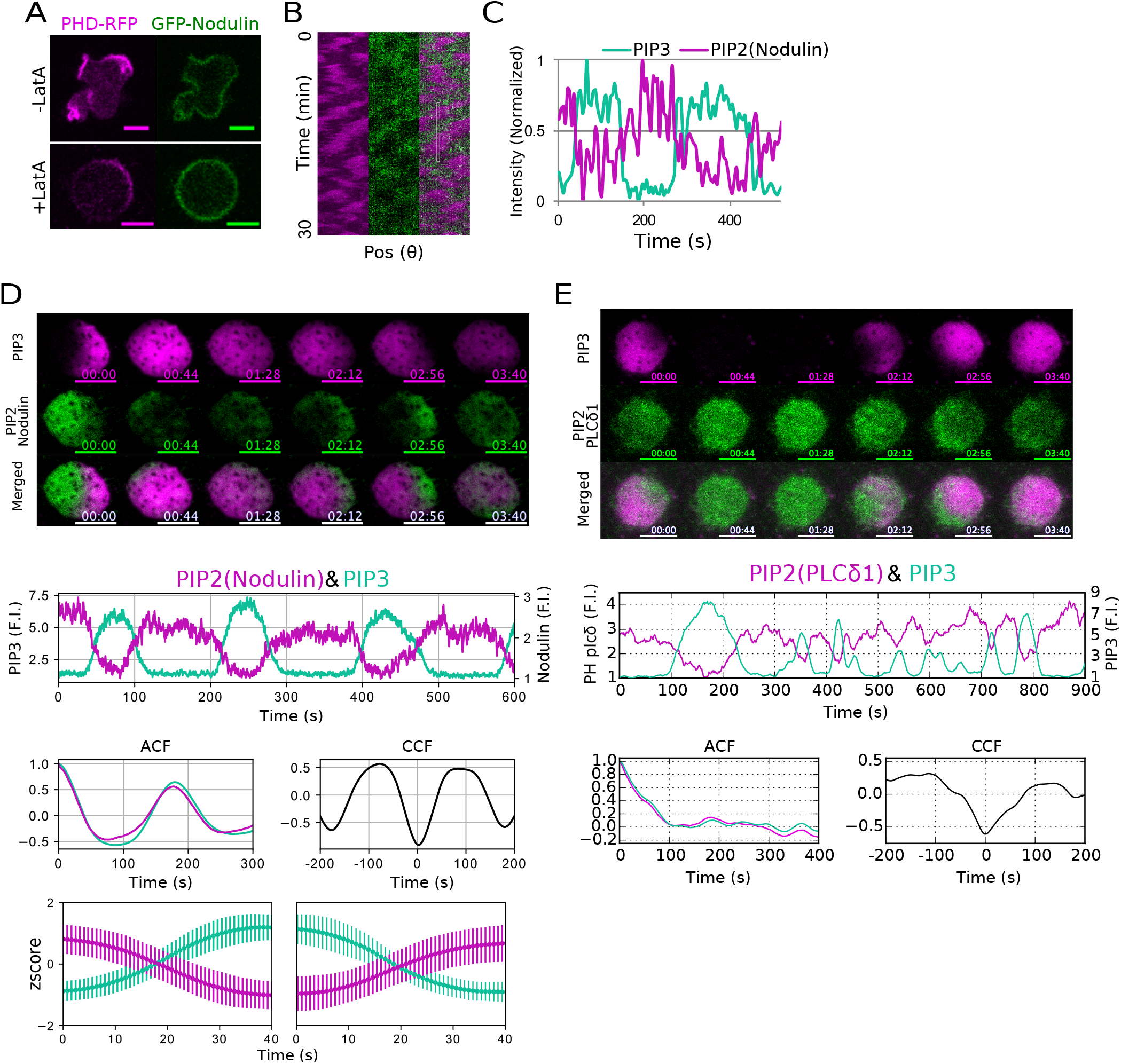
Observation of PIP2 with Nodulin and PHD_PLCδ1_. (A) Confocal images of PIP3 (PHD_AKT/PKB_-RFP) and GFP-Nodulin before and after treatment with Latrunculin A. (B) Kymographs of the cell periphery shown in S7A Fig, bottom. (C) Time trajectory of PIP3 and PIP2 localization patterns in the ROI shown in the kymograph. (D), (E) Simultaneous time-lapse TIRFM imaging, typical time trajectories, auto- and cross-correlation functions and average trajectories of simultaneous TIRF time-lapse imaging of PIP3 and GFP-Nodulin and PHD_PLCδ1_-GFP. Scale bars, 5 *μ*m. Time format is “mm:ss”.

**S8 Fig. S8.**
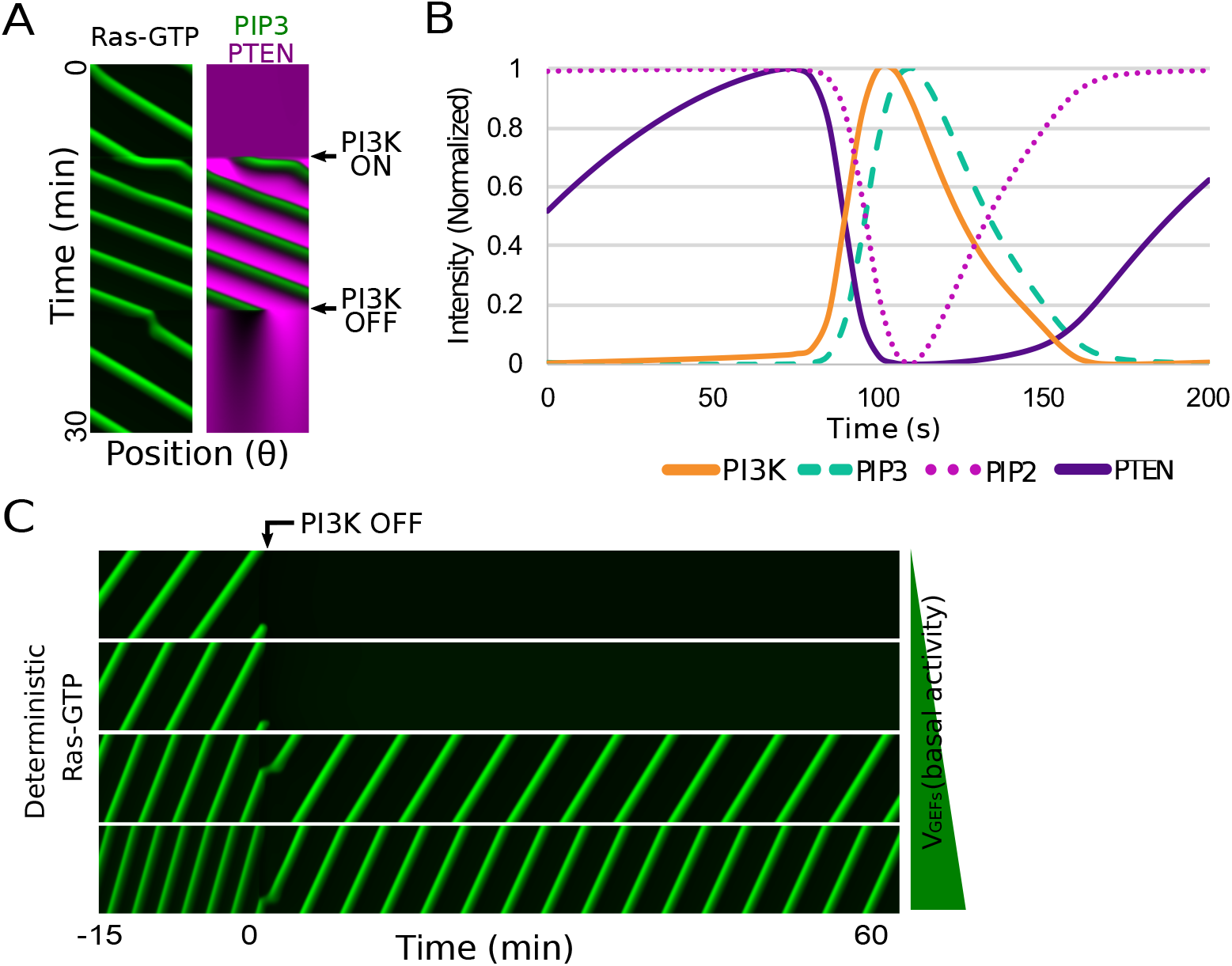
Deterministic simulation of Ras-GTP wave. (A) The kymograph generated by a simulation based on the model. V_GEFs_ = 550 s^-1^. (B) Time trajectory of Ras-GTP/PI3K, PIP3, PIP2 and PTEN obtained from the simulation. (C) Simulation results of the PI3K inhibition experiment. The termination of Ras waves depended on the basal activity (V_GEFs_ = 450, 500, 550, 600 s^-1^).

**S1 Table.** Parameters for simulation (PIP3).

| Parameters | Description | Values |
| --- | --- | --- |
| $V_{PTEN}$ | Dephosphorylation rate of PIP3 by PTEN | $3 \text{ s}^{-1}$ |
| $K_{PTEN}$ | Michaelis constant of PTEN dephosphorylation reaction | $500 \mu\text{m}^{-2}$ |
| $V_{PI3K}$ | Phosphorylation rate of PIP2 by PI3K | $12 \text{ s}^{-1}$ |
| $K_{PI3K}$ | Michaelis constant of PI3K phosphorylation reaction | $300 \mu\text{m}^{-2}$ |
| $V_{PTEN\_ass}$ | PTEN association reaction rate by PIP2 | $5000 \text{ molecules} \cdot \mu\text{M}^{-1} \cdot \mu\text{m}^{-2} \cdot \text{s}^{-1}$ |
| $K_{PIP2}$ | Michaelis constant of PTEN association reaction | $2000 \mu\text{m}^{-2}$ |
| $\lambda_{PIP3}$ | PTEN-independent PIP3 degradation rate | $0.2 \text{ s}^{-1}$ |
| $\beta$ | Parameter indicates magnitude of PI3K activation by Ras | 0.01 |
| $\chi$ | A constant to transform surface concentration to volume concentration | $0.001 \text{ molecules} \cdot \mu\text{M} \cdot \mu\text{m}^{-2}$ |
| $PTEN_{total}$ | Total concentration of PTEN | $0.1 \mu\text{M}$ |
| $PIP_{total}$ | Density of PIPs on plasma membrane | $5000 \text{ molecules} \cdot \mu\text{m}^{-2}$ |
| $R$ | Cell radius | $5 \mu\text{m}$ |
| $D$ | Diffusion coefficient of molecules on plasma membrane | $0.2 \mu\text{m}^{-2} \cdot \text{s}^{-1}$ |
| $[PIP3]_{typical}$ | Typical value of [PIP3] | $\sim 200 \mu\text{m}^{-2}$ |
| $[PTEN]_{typical}$ | Typical value of [PTEN] | $\sim 400 \mu\text{m}^{-2}$ |

**S2 Table.**
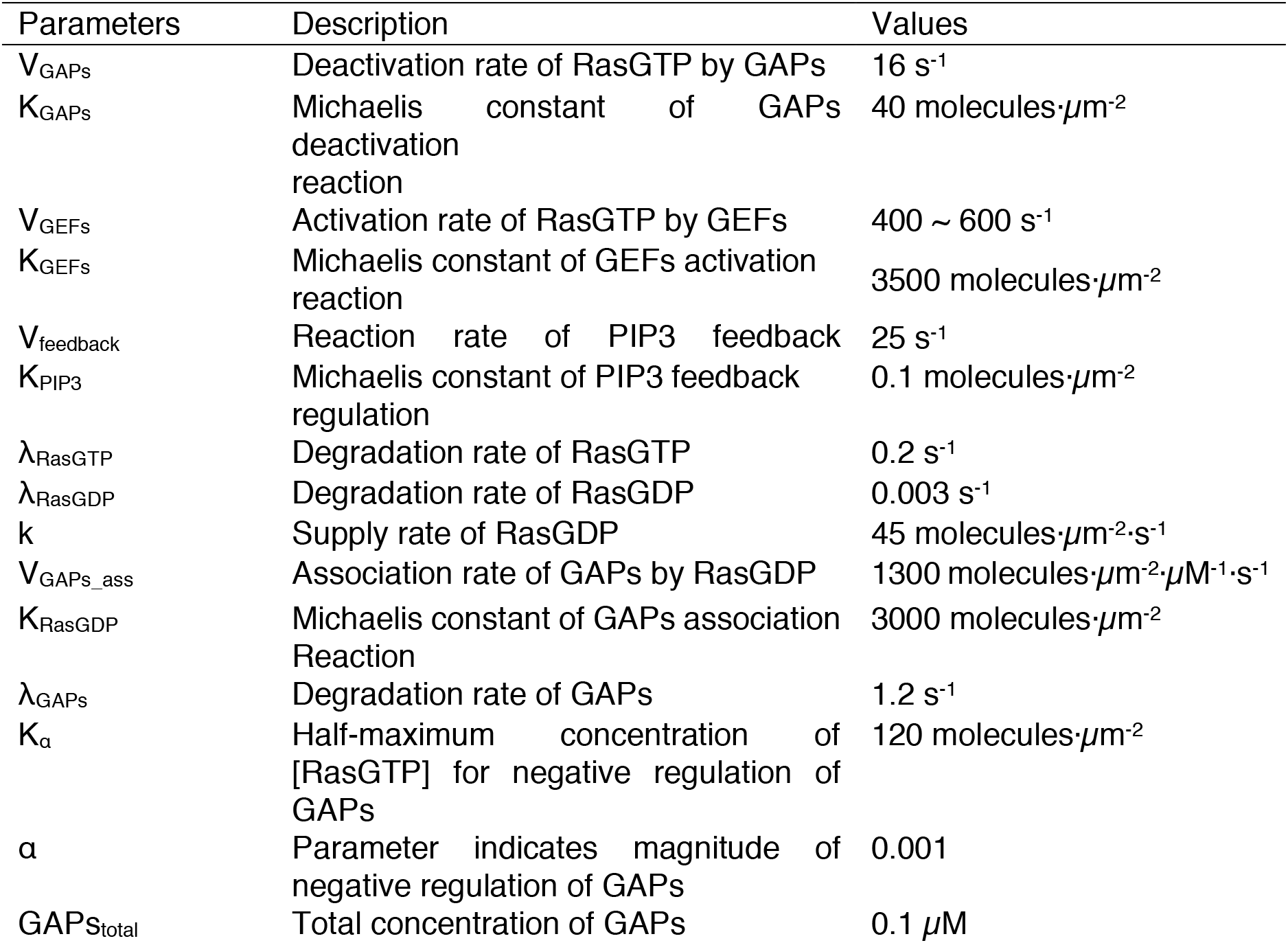
Parameters for simulation (Ras).

| Parameters | Description | Values |
| --- | --- | --- |
| $V_{\text{GAPs}}$ | Deactivation rate of RasGTP by GAPs | $16 \text{ s}^{-1}$ |
| $K_{\text{GAPs}}$ | Michaelis constant of GAPs deactivation reaction | $40 \text{ molecules} \cdot \mu\text{m}^{-2}$ |
| $V_{\text{GEFs}}$ | Activation rate of RasGTP by GEFs | $400 \sim 600 \text{ s}^{-1}$ |
| $K_{\text{GEFs}}$ | Michaelis constant of GEFs activation reaction | $3500 \text{ molecules} \cdot \mu\text{m}^{-2}$ |
| $V_{\text{feedback}}$ | Reaction rate of PIP3 feedback | $25 \text{ s}^{-1}$ |
| $K_{\text{PIP3}}$ | Michaelis constant of PIP3 feedback regulation | $0.1 \text{ molecules} \cdot \mu\text{m}^{-2}$ |
| $\lambda_{\text{RasGTP}}$ | Degradation rate of RasGTP | $0.2 \text{ s}^{-1}$ |
| $\lambda_{\text{RasGDP}}$ | Degradation rate of RasGDP | $0.003 \text{ s}^{-1}$ |
| $k$ | Supply rate of RasGDP | $45 \text{ molecules} \cdot \mu\text{m}^{-2} \cdot \text{s}^{-1}$ |
| $V_{\text{GAPs\_ass}}$ | Association rate of GAPs by RasGDP | $1300 \text{ molecules} \cdot \mu\text{m}^{-2} \cdot \mu\text{M}^{-1} \cdot \text{s}^{-1}$ |
| $K_{\text{RasGDP}}$ | Michaelis constant of GAPs association Reaction | $3000 \text{ molecules} \cdot \mu\text{m}^{-2}$ |
| $\lambda_{\text{GAPs}}$ | Degradation rate of GAPs | $1.2 \text{ s}^{-1}$ |
| $K_{\alpha}$ | Half-maximum concentration of [RasGTP] for negative regulation of GAPs | $120 \text{ molecules} \cdot \mu\text{m}^{-2}$ |
| $\alpha$ | Parameter indicates magnitude of negative regulation of GAPs | 0.001 |
| $\text{GAPs}_{\text{total}}$ | Total concentration of GAPs | $0.1 \mu\text{M}$ |
| $[\text{RasGTP}]_{\text{typical}}$ | Typical value of [RasGTP] | $\sim 700 \mu\text{m}^{-2}$ |
| $[\text{RasGDP}]_{\text{typical}}$ | Typical value of [RasGDP] | $\sim 4000 \mu\text{m}^{-2}$ |
| $[\text{GAPs}]_{\text{typical}}$ | Typical value of [GAPs] | $\sim 50 \mu\text{m}^{-2}$ |

**S1 Movie. RBD_Raf1_-RFP & PHD_AKT/PKB_-GFP / Ax2.** Time-lapse confocal images show RBD_Raf1_-RFP (left), PHD_AKT/PKB_-GFP (center) and merged (right), corresponding to Fig 1A. Scale bars represent 5 μm. Time format is “mm:ss”.

**S2 Movie. RBD_Raf1_-RFP & PHD_AKT/PKB_-GFP / Ax2 (TIRF).** Time-lapse TIRF images show RBD_Raf1_-RFP (left), PHD_AKT/PKB_-GFP (center) and merged (right), corresponding to Fig 2A. Scale bars represent 10 μm. Time format is “mm:ss”.

**S3 Movie. PI3K2-Halo-TMR & RBD_Raf1_-GFP/Ax2 (TIRF).** Time-lapse TIRF images show PI3K2-Halo-TMR (left), RBD_Raf1_-GFP (center) and merged (right), corresponding to S2B Fig.

**S4 Movie. PI3K2-Halo-TMR & PHD_AKT/PKB_-GFP/Ax2 (TIRF).** Time-lapse TIRF images show PI3K2-Halo-TMR (left), PHD_AKT/PKB_-GFP (center) and merged (right), corresponding to S3A Fig.

**S5 Movie. PI3K2(KE)-Halo-TMR & RBD_Raf1_-GFP/Ax2 (TIRF).** Time-lapse TIRF images show PI3K2^K857, 858E^-Halo-TMR (left), RBD_Raf1_-GFP (center) and merged (right), corresponding to Fig 3A.

**S6 Movie. PHD_AKT/PKB_-RFP & GFP-Nodulin/Ax2 (TIRF).** Time-lapse TIRF images show PHD_AKT/PKB_-RFP (left), GFP-Nodulin (center) and merged (right), corresponding to S7D Fig.

**S7 Movie. PTEN-Halo-TMR & PHD_AKT/PKB_-GFP /Ax2, PTEN-Halo-TMR & GFP-Nodulin /Ax2, RFP-Nodulin & RBD_Raf1_-GFP /Ax2, PI3K2-Halo-TMR & GFP-Nodulin /Ax2, PTEN-Halo-TMR & RBD_Raf1_-GFP /Ax2, PI3K2-Halo-TMR & PTEN-GFP /Ax2 (TIRF).** Time-lapse TIRF images show RFP or Halo-TMR conjugated proteins (left), GFP conjugated proteins (center) and merged (right), corresponding to S5 and S6 Fig.

**S8 Movie. RBD_Raf1_-GFP (+100 *μ*M LY294002).** Time-lapse confocal images show RBDRaf1-GFP with 100 *μ*M LY294002, corresponding to Fig 4A. LY294002 was added at time 1 min. Scale bars represent 20 μm. Time format is “mm:ss”.

